# Targeting GL-Lect driven endocytosis to suppress cell plasticity in breast cancer

**DOI:** 10.64898/2026.01.08.698324

**Authors:** Amir Fardin, Andrea Francesco Benvenuto, Irene Schiano Lomoriello, Chiara Tordonato, Micaela Quarto, Stefano Freddi, Giovanni Giangreco, Giovanni Bertalot, Francesca Montani, Ivan Nicola Colaluca, Roberta Cacciatore, Andrea Raimondi, Maria Giovanna Jodice, Maria Grazia Malabarba, Luigi Scietti, Giuseppe Ciossani, Alessandro Cuomo, Massiullah Shafaq-Zadah, Estelle Dransart, Sina Kanannejad, Brenda Green, Salvatore Pece, Stefano Confalonieri, Ulf J. Nilsson, Hakon Leffler, Giorgio Scita, Ludger Johannes, Pier Paolo Di Fiore, Sara Sigismund

## Abstract

Aberrant endocytosis has long been associated with epithelial plasticity and tumorigenesis, but direct *in vivo* evidence of its causal role in tumor progression and metastasis has been lacking. Here, we identify and molecularly characterize a previously unrecognized form of E-cadherin (ECAD) internalization in mammary epithelial cells. This process is mediated by the endocytic adaptor Epsin 3 (EPN3) through glycolipid-lectin (GL-Lect) driven endocytosis requiring galectin-3 and Eps15-family adaptors. Leveraging an EPN3 knock-in mouse model, we show that dysregulation of GL-Lect driven endocytosis disrupts mammary gland morphogenesis and activates epithelial-to-mesenchymal plasticity (EMP), synergizing with the ERBB2/Neu breast oncogene to drive metastasis. Pharmacologic inhibition of the GL-Lect mechanism suppresses morphogenetic and invasive phenotypes *ex vivo*, providing proof-of-concept for therapeutic targeting. These findings establish the GL-Lect mechanism as a driver of metastatic plasticity and uncover a tractable vulnerability in BC.

## Introduction

Epithelial-to-mesenchymal transition (EMT) is a dynamic process involving transcriptional and phenotypic changes that alter cell polarity, adhesion, and motility. It plays essential roles in development and tissue remodeling and is a key driver of invasiveness and metastasis in cancer^1, 2^. Successful metastatic colonization, however, requires the reverse mesenchymal-to-epithelial transition (MET) to re-establish epithelial traits at distant sites^3, 4^.

EMT is not a binary event but rather a continuum of intermediate or hybrid states - termed epithelial-to-mesenchymal plasticity (EMP) - characterized by co-expression of epithelial and mesenchymal markers ^5^. Cells in these metastable states exhibit enhanced plasticity, enabling dynamic switching between invasive and proliferative phenotypes. In the case of the mammary gland, this plasticity is important for both physiological processes, such as ductal branching and involution^2^, and pathological events such as breast cancer (BC) metastasis^5–7^.

Recent evidence links EMP to the regulation of epithelial markers such as E-cadherin (ECAD) through endocytic trafficking rather than solely transcriptional repression^8, 9^. By modulating endocytic pathways and intracellular trafficking, cells can reversibly transition between phenotypic states with greater flexibility. In this context, we identified the endocytic protein Epsin 3 (EPN3) as a driver of EMP in the normal-like breast epithelial cell line MCF10A^10^. EPN3 is a member of the epsin family of endocytic adaptor proteins, which also includes Epsin 1 (EPN1) and Epsin 2 (EPN2)^11, 12^. EPN3 overexpression promotes ECAD endocytosis, disrupts cell-cell junctions, activates β-catenin signaling, induces mesenchymal gene expression, and establishes a TGFβ autocrine loop, collectively enhancing cancer stem cell traits and invasiveness. These effects are specific to EPN3 and are not replicated by its paralog EPN1^10^.

EPN3 is overexpressed in approximately 50% of BCs, with 10% harboring gene amplification, and half of these cases exhibiting co-amplification of the BC oncogene ERBB2 (HER2). EPN3 is overexpressed in approximately 50% of BCs, with 10% harboring gene amplification, and half of these cases exhibiting co-amplification of the BC oncogene ERBB2 (HER2). Thus, EPN3 is one of the few endocytic adaptor proteins exhibiting genetic alterations, which can therefore be unambiguously detected in tumors. Clinically, high EPN3 expression correlates with poor prognosis and increased distant metastasis risk independent of ERBB2 status^10^. Here, we delineate an EPN3-mediated glycolipid-lectin (GL-Lect) driven endocytic process involving galectin-3 (Gal3) and glycosphingolipids (GSLs), which orchestrates ECAD internalization and phenotypic plasticity. We further characterize molecular determinants within EPN3 that confer its specificity in driving ECAD endocytosis, and we reveal a key role for Eps15 family adaptors in ECAD clustering and Gal3 recruitment. Importantly, we identify inhibitors targeting this endocytic mechanism, highlighting its therapeutic potential.

To validate these findings *in vivo*, we generated an epithelial-specific EPN3 knock-in (EPN3-KI) mouse model, in which EPN3 is overexpressed at levels comparable to those detected in human BCs. These mice exhibit aberrant mammary gland morphogenesis, including enhanced branching and alveologenesis. Single-cell RNA sequencing (scRNA-seq) revealed upregulation of EMP in basal cells, and metabolic and lactation-related programs in luminal cells. Inhibition of GL-Lect driven ECAD endocytosis reversed these phenotypes in primary organoids, linking EPN3 function to mammary tissue remodeling. Moreover, EPN3 overexpression accelerated tumorigenesis and lung metastasis in MMTV-NeuN mice, underscoring its role in BC progression.

Collectively, this study establishes endocytosis as a critical regulator of EMP and BC metastasis. Given the prevalence of EPN3 overexpression in BC^10^, targeting its associated endocytic processes represents a promising strategy to impede tumor invasiveness and dissemination.

## Results

### EPN3 promotes clathrin-independent ECAD endocytosis via the GL-Lect mechanism and Eps15 family proteins

We previously demonstrated that accelerated ECAD endocytosis is the initiating event in a cascade triggered by EPN3 overexpression ultimately leading to EMP in the non-transformed breast epithelial cell line MCF10A cells^10^. To elucidate the endocytic mechanism underlying EPN3-induced ECAD internalization, we first examined its dependence on clathrin-mediated endocytosis (CME). Since clathrin knockdown (KD) is highly cytotoxic due to its multiple roles in the cell beyond internalization, such as in Golgi trafficking and mitosis^13^, we silenced the major clathrin adaptor AP2. Immunofluorescence (IF)-based internalization assays following AP2 KD revealed no impact on EPN3-driven ECAD endocytosis, whereas internalization of the transferrin receptor (TfR), a canonical CME cargo, was markedly impaired (**Fig. 1A and S1A**).

**Figure 1.**
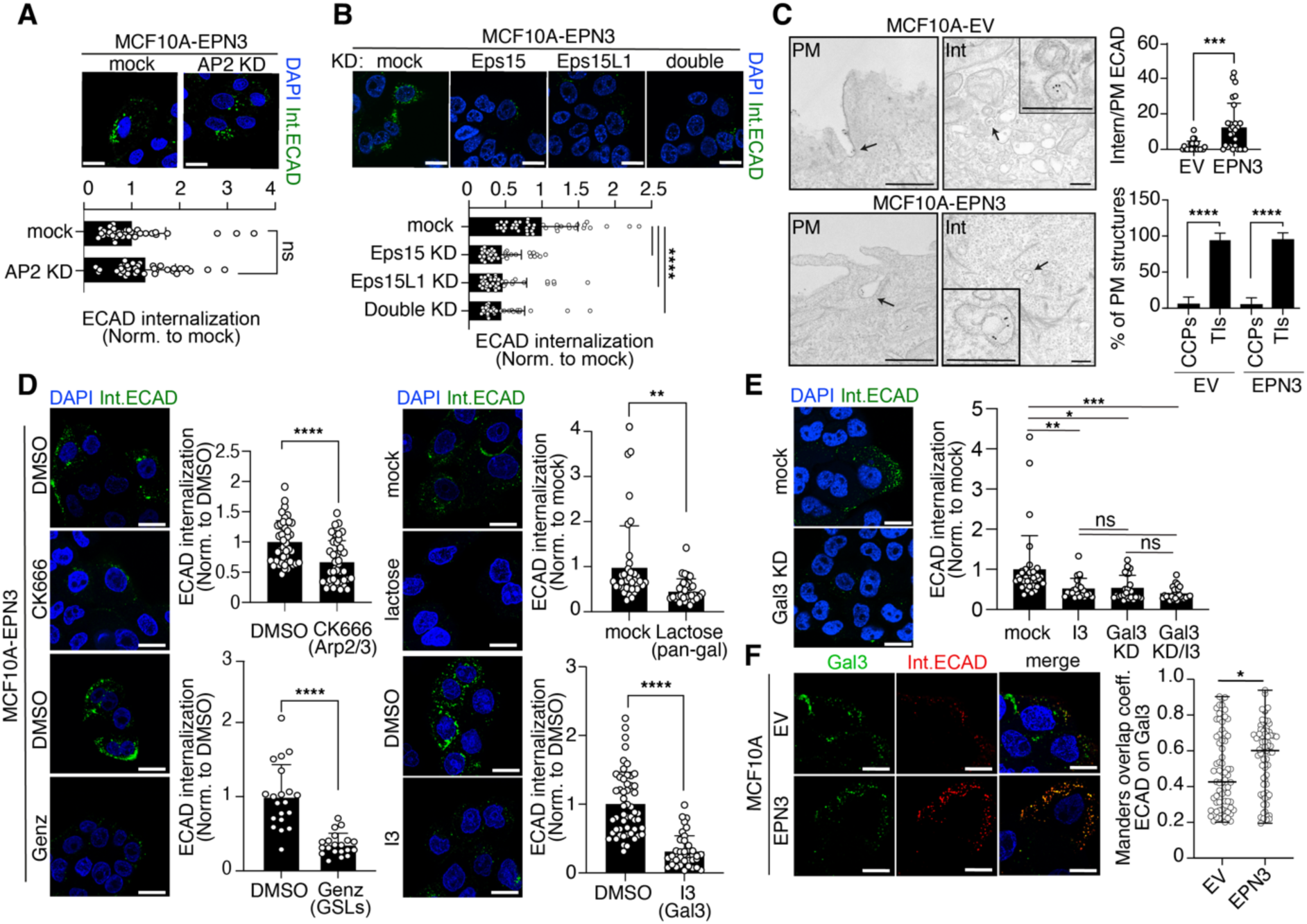
EPN3-mediated ECAD endocytosis in MCF10A cells. **A)** ECAD internalization was monitored by IF in MCF10A-EPN3 cells treated with AP2µ KD or mock. Top, representative images; internalized ECAD (green), DAPI (blue). Bar, 20 µm. Bottom, quantification of relative internalized ECAD fluorescence intensity/cell in individual field of views, normalized to mock control. N (fields of view): Mock=4, AP2µ=35; n=3. **B)** ECAD internalization in MCF10A-EPN3 cells subjected to single or double Eps15/Eps15L1 KDs. Representative images and quantification as in (A). Bar, 20 µm. N (fields of view): Mock=42, Eps15 KD=39, Eps15L1 KD=40, Double KD=35, n=3. **C)** Left panels: PM, representative immuno-EM images showing PM-ECAD-positive (gold-labeled) tubular invaginations (indicated by arrows) in MCF10A-EV and MCF10A-EPN3 cells; scale bar, 200 nm; Int, representative immuno-EM images showing internalized ECAD-positive (gold-labeled) structures (indicated by arrows and enlarged in the insets) in MCF10A-EV and MCF10A-EPN3 cells; scale bar, 250 nm. Right upper panel: quantification of internalized ECAD expressed as a percentage of PM-ECAD. N (cells): EV=24, EPN3=27. Right-lower panel: gold-labelled ECAD-positive clathrin-coated pits (CCPs) and tubular invaginations (TIs) expressed as percentage of total number of structures in 100 µm PM length/cell. N (cells): EV=27, EPN3=25. **D)** Effects of inhibitors on ECAD internalization in MCF10A-EPN3 cells. Cells were pre-treated with the indicated compounds or vehicle (DMSO) before measuring ECAD internalization as in (A): CK666 (50 µM, 1 h), Genz-123346 (4 µM, 6 days), Lactose (100 mM, 1h), I3 (20 µM, 10 min). Representative images and quantification as in (A). Bar, 20 µm. N (fields of view): CK666=42 (DMSO control=44) (n=5); Genz=20 (DMSO control=20) (n=3); Lactose=33 (mock=40) (n=3); I3=40 (DMSO control=53) (n=6). **E)** ECAD internalization in MCF10A-EPN3 cells subjected to I3 treatment as in (D), Gal3 KD or Gal3 KD/I3 treatment. Representative images and quantification as in (A). Bar, 20 µm. N (fields of view): Mock=34, I3=23, Gal3 KD=26, Gal3 KD/I3=28, n=2. **F)** Co-internalization of Gal3–ECAD was monitored for 10 min in MCF10A-EV and -EPN3 cells. Left: representative confocal images, Gal3-Alexa488 (green), anti-ECAD (red), DAPI (blue). Bar: 20 µm. Right: Manders overlap coefficient of internalized Gal3-ECAD. N (cells): EV/EPN3=69; n=2. In all panels, results are shown as mean±SD, except panel for E in which median ± max/min values are shown. p-values (unpaired Student’s t-test, two-tailed): **** <0.0001; *** <0.001; ** <0.01; * <0.05, ns, not significant.

The endocytic adaptors Eps15 and Eps15L1 have been implicated - together with epsins - in both clathrin-dependent and clathrin-independent endocytic mechanisms^14, 15^. To assess their role in EPN3-driven ECAD internalization, we performed single and double KD experiments in EPN3-overexpressing MCF10A cells (MCF10A-EPN3). Silencing of Eps15 or Eps15L1 individually significantly impaired ECAD endocytosis, while their combined KD did not result in further inhibition, indicating essential, non-redundant roles for these proteins (**Fig. 1B and S1B**). These findings suggest that a clathrin-independent, Eps15/Eps15L1-dependent mechanism is involved in EPN3-driven ECAD endocytosis.

To further confirm the clathrin-independent nature of this process, we analyzed the ECAD-internalizing structures by electron microscopy (EM). MCF10A-EPN3 cells displayed elevated levels of ECAD internalization compared to empty vector controls (MCF10A-EV) (**Fig. 1C, top right**). In both cell lines, the majority of plasma membrane (PM) ECAD localized to uncoated tubular invaginations (TIs), consistent with a clathrin-independent route, while only a negligible fraction was found in clathrin-coated pits (CCPs, **Fig. 1C, bottom right**). Internalized ECAD was often found in crescent-shaped short tubular structures (**Fig. 1C, Int**) that had the typical morphology of clathrin-independent carriers^16^.

To further characterize this process, we conducted an inhibitor screening of various endocytic mechanisms (see **Fig. 1D and Fig. S1C-E,** for positive and negative controls). Inhibition of actin polymerization (via the Arp2/3 inhibitor CK666), glycosphingolipid (GSL) synthesis (via long-term treatment with Genz-123346) or galectin-glycan interactions (via the non-selective pan-galectin inhibitor lactose) all significantly reduced ECAD endocytosis (**Fig. 1D)**. Notably, the same effect was observed following acute inhibition (10 min) of Gal3 using the potent, selective, non-membrane-permeable inhibitor I3 (K_d_ of 99 ± 4 nM and 120 ± 21 nM for human and mouse Gal3, respectively, see Methods)^17^ (**Fig. 1D, bottom right)** or upon Gal3 knockdown (KD) (**Fig. 1E, S1F,G** for positive and negative controls). Importantly, the combination of I3 treatment and Gal3 KD did not further reduce E-cadherin (ECAD) endocytosis (**Fig. 1E**), confirming the specificity of the inhibitor. In contrast, inhibitors/KD targeting micropinocytosis^18^, caveolae-mediated endocytosis^19^ and EGFR non-clathrin endocytosis (EGFR-NCE)^20^ had no effect (**Table S1A**). To further confirm the involvement of Gal3, we performed live-cell imaging of Alexa488-labeled Gal3 and ECAD. In control cells, ECAD-positive internalizing vesicles exhibited partial colocalization with Alexa488-Gal3 after 10 minutes of endocytosis. This colocalization was markedly increased in MCF10A-EPN3 cells (**Fig. 1F**), consistent with the enhanced Gal3-dependent ECAD endocytosis observed in these cells.

We note that ECAD has previously been shown to undergo internalization through both clathrin-dependent and clathrin-independent mechanisms, depending on the context, with clathrin-independent endocytosis observed in breast cancer cell lines^21^. Our findings show that, in mammary epithelial cells, EPN3 overexpression induces ECAD endocytosis via a clathrin-independent mechanism that exhibits the characteristics of GL-Lect driven endocytosis^22^, relying on GSLs and Gal3-glycan interactions. Moreover, this process requires the involvement of the Eps15/Eps15L1 protein family, which has not previously been implicated in this endocytic route. Notably, the same mechanism operates in MCF10A control cells, albeit with lower magnitude (**Fig. S1H-N**), indicating that EPN3 overexpression amplifies an existing physiological process. These observations are consistent with our previous findings that EPN3 depletion reduces basal ECAD endocytosis in MCF10A cells^10^.

### Molecular basis of EPN3 specificity in ECAD endocytosis

We previously demonstrated that EPN3 overexpression in MCF10A cells promotes EMP and enhances cellular invasion - phenotypes not recapitulated by EPN1 overexpression^10^. Notably, in contrast to EPN3, EPN1 neither induces ECAD endocytosis (**Fig. 2A, top**) nor morphological changes (elongation) characteristic of EMT (**Fig. 2A, bottom**).

**Figure 2.**
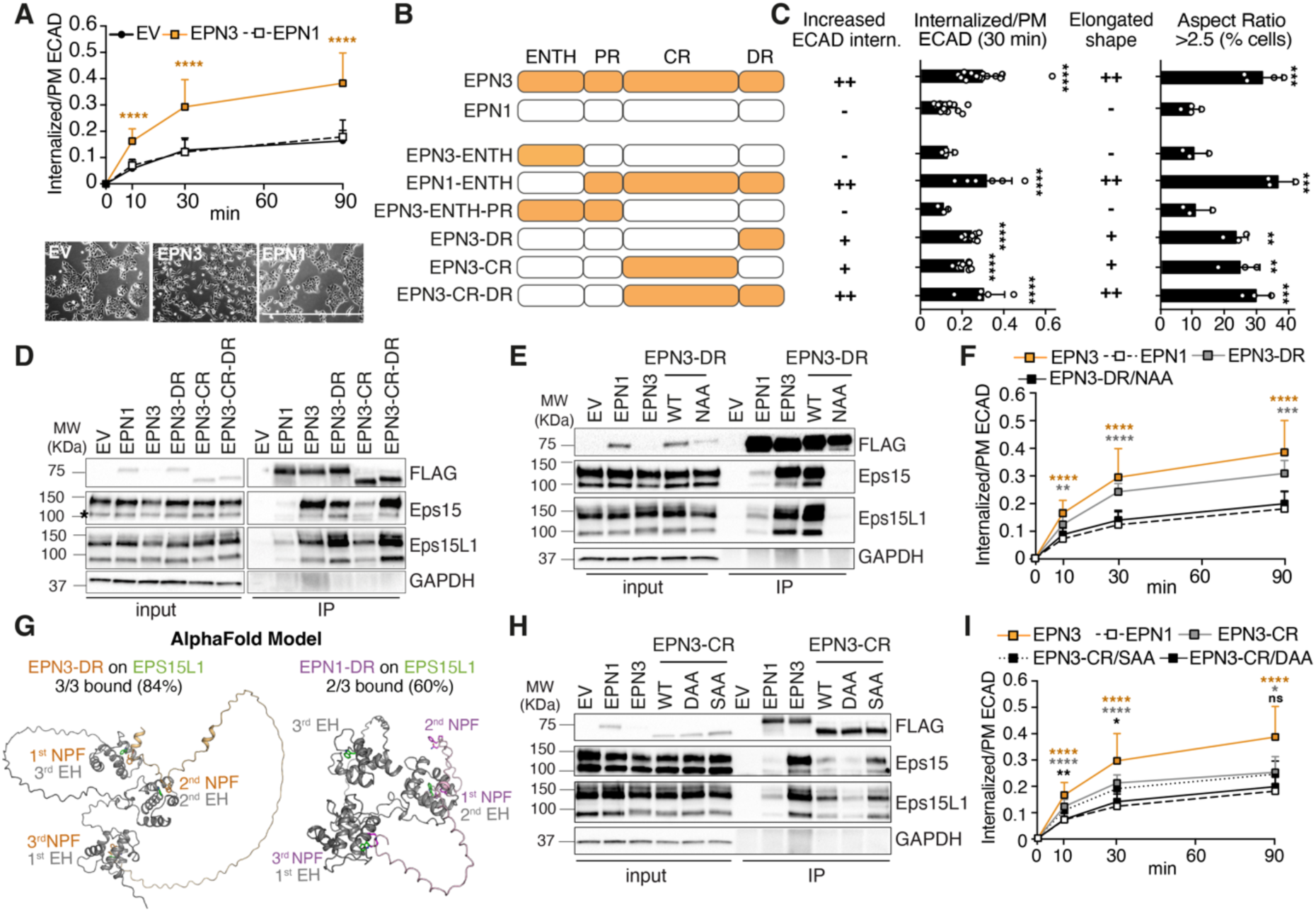
Structure-function analysis of EPN1 and EPN3. **A)** Top, Time-course of ECAD internalization measured by FACS in MCF10A cells stably transfected with constructs expressing doxycycline (doxy)-inducible FLAG-EPN3 or -EPN1, or empty vector (EV). Expression was induced for 24 h. Data are reported as mean fluorescence intensity of internalized ECAD signal over total plasma membrane (PM) signal ± SD (n≥4). Bottom, representative phase contrast microscopy images. Bar, 1000 μm. **B)** Schematics of the FLAG-tagged EPN1-EPN3 chimeras, expressed as doxy-inducible constructs in MCF10A cells. Four domains are highlighted: epsin-N-terminal homology domain (ENTH), proximal region (PR), central region (CR) and distal region (DR). Details are in **Fig. S2**. **C)** Left, summary of ECAD internalization measured by FACS (actual data are in **Fig. S3B**) for the constructs shown in B: ++, maximal endocytosis (comparable to EPN3); +, intermediate endocytosis; -, minimal endocytosis (comparable to EPN1). Middle left, the 30 min time point of internalized ECAD/PM (**Fig. S3B**) is shown ±SD; n≥3. Middle right, summary of the elongated morphology of MCF10A cells expressing the indicated constructs (scores ++, + e – are as from the left panel). Right, phalloidin staining (details are in **Fig. S3C**) was used to measure the aspect ratio of each cell. Percentage of cells with an aspect ratio > 2.5 is shown ± SD; n≥3. **E)** Co-immunoprecipitation of the indicated EPN1/EPN3 chimeras with Eps15/Eps15L1. Anti-FLAG immunoprecipitates (IP) from cells as in A-C were immunoblotted (IB) as shown on the right. Inputs and IPs were run on the same gel/IB. Asterisk indicates a non-specific band in the anti-Eps15 IB (see also **Table S1B** for IB in Eps15 KD cells). **F)** Role of NPF motifs. Anti-FLAG IP from the indicated transfectants were IB as indicated on the right. **G)** Time-course of ECAD internalization measured by FACS in MCF10A cells expressing the indicated constructs. Data are reported as mean fluorescence intensity of internalized ECAD signal over the PM signal at time=0 ± SD; n≥5. **H)** AlphaFold3 predictions of the complex between the EPN1-DR or EPN3-DR regions and the Eps15L1 EH domains. For EPN3-DR, 84% of the generated models (n=50), showed all three NPFs binding to the three EPS15L1 EH domains. For EPN1-DR, 60% of the generated models (n=50) showed two NPFs engaging two Eps15L1 EH domains. Additional models and relative predicted aligned error (PAE) plots for each binding model (including models with Eps15) are shown in **Fig. S5**. **I)** Role of DPF/SWG motifs. Anti-FLAG IP from the indicated transfectants were IB as indicated on the right. **L)** Time-course of ECAD internalization measured by FACS of MCF10A cells expressing the indicated constructs. Data are reported as mean fluorescence intensity of internalized ECAD signal over the PM signal at time=0 ± SD; n≥6. p-values are as in Fig. 1. In the various IB, GAPDH is used as negative control for IP and loading control for inputs and MWs are shown on the left in KDa.

Given the high sequence homology and conserved domain organization between EPN3 and EPN1 (**Fig. S2**), we generated EPN3-EPN1 chimeric proteins (**Fig. 2B** and **Fig. S3A**), expressed in MCF10A cells under a doxycycline-inducible promoter, to identify the molecular determinants responsible for EPN3-specific phenotypes (ECAD internalization and elongation).

Swapping the ENTH domains, which share 80% identity and 95% similarity (**Fig. S2**), did not alter EPN3 function nor confer EPN3-like phenotypes to EPN1, suggesting that this domain does not mediate EPN3-specific activities (**Fig. 2B,C** and **Fig. S3B-D**). Similarly, the EPN3 proximal region (PR), containing ubiquitin-binding motifs (UIMs) (**Fig. S2**), did not account for the functional specificity of EPN3. This was evidenced by the EPN3-ENTH-PR chimera, composed of the ENTH and PR domains of EPN3 and the remaining regions from EPN1, which failed to reproduce EPN3-like phenotypes (**Fig. 2B,C** and **Fig. S3B-D**).

In contrast, the C-terminal region, which exhibits the highest divergence between the two proteins, appears to contain key determinants of EPN3-specific activity. Chimeras incorporating either the central region (CR), which includes DPF/DPW motifs and putative clathrin-binding sites, or the distal region (DR), enriched in NPF motifs (EPN3-CR and EPN3-DR, respectively; **Fig. S2**), exhibited partial phenotypic rescue, with EPN3-DR displaying a higher endocytic rate (**Fig. S3B-D**). Of note, the simultaneous substitution of both regions (EPN3-CR-DR) fully restored the EPN3-specific phenotype (**Fig. 2B,C** and **Fig. S3B-D**). These results indicate that the CR and DR regions both contribute to the unique functional properties of EPN3 in ECAD endocytosis.

### A dual Eps15/Eps15L1-binding surface in EPN3 mediates ECAD endocytosis

To elucidate the molecular basis by which the CR and DR regions of EPN3 confer functional specificity relative to EPN1, we evaluated the interaction of wild-type (WT) and chimeric EPN3 and EPN1 proteins with Eps15 family members. Co-immunoprecipitation (co-IP) assays revealed that WT EPN3 exhibits significantly stronger binding to Eps15 and Eps15L1 compared to EPN1 (**Fig. 2D**), consistent with an Eps15/Eps15L1-dependent mechanism underlying ECAD endocytosis.

Notably, the EPN3-DR chimera exhibited an almost complete, though not full, rescue of binding to Eps15 and Eps15L1, whereas the EPN3-CR chimera showed a milder but consistent rescue (**Fig. 2D**), mirroring their respective effects on ECAD endocytosis (**Fig. S3B-D**). Importantly, both regions are required for optimal binding to Eps15 and Eps15L1, as evidenced by the restored full binding capacity of the EPN3-CR-DR chimera (**Fig. 2D).** Together, these findings demonstrate that the CR and DR domains of EPN3 provide distinct Eps15/Eps15L1-binding surfaces both of which contribute to drive the assembly of EPN3-Eps15/Eps15L1 complexes.

Within the DR domain, three NPF motifs (**Fig. S2B**) - known mediators of interactions with EH-domain-containing proteins such as Eps15 family members^23^ – were identified as candidate binding sites. Site-directed mutagenesis of these motifs to NAA in the EPN3-DR chimera (**Fig. S4A**) abolished binding to both Eps15 and Eps15L1 (**Fig. 2E**) and abrogated EPN3-specific phenotypes (**Fig. 2F** and **Fig. S4B,C**), confirming the essential role of NPF motifs in the interaction. Intriguingly, despite harboring three NPF motifs (**Fig. S2B**), EPN1 exhibited weak binding to Eps15/Eps15L1, suggesting that differences in the three-dimensional conformation of their DR domains may underlie the differential binding affinities.

To investigate this, we performed AlphaFold3 (AF3) modelling of the complex between the EPN3-DR or EPN1-DR domains and the three EH domains of Eps15 or Eps15L1. In all the generated models, at least two out of three NPF motifs in both EPN1 and EPN3 bound to the Eps15 or EpsL1 EH domains. Interestingly, the most probable configuration for the EPN3-Eps15L1 interaction involved three NPFs bound to three EH domains (**Fig. 2G**, 84% of the cases). In contrast the preferred binding configuration for EPN1-Eps15L1 involved two NPFs and two EHs (**Fig. 2G**, 60% of the cases). Similar observations were made for the interactions of EPN3-DR or EPN1-DR with Eps15 EH domains (**Fig. S5**, where further analyses and statistics are also shown). These data parallel the experimental observation and further suggest that the overall binding propensity of EPN3 for the EH domains of Eps15/is higher than that of EPN1.

Analysis of the CR domain revealed EPN3-specific sequence motifs - a SWG motif and a DPF motif (**Fig. S2B**) – both conserved in human and mouse EPN3 and previously reported as non-canonical EH-domain binding sites^24–26^. In the EPN3-CR chimera, mutation of the DPF motif to DAA significantly impaired binding to Eps15/Eps15L1 and inhibited ECAD endocytosis, whereas mutation of the SWG motif to SAA had no detectable effects (**Fig. 2H,I; Fig. S4**).

Collectively, these data identify a dual Eps15/Eps15L1-binding surface in EPN3, comprised of a DPF motif within the CR domain and three NPF motifs within the DR domain, which together confer the protein’s functional specificity in mediating ECAD endocytosis.

### EPN3-Eps15/Eps15L1 complexes facilitate ECAD clustering and Gal3 recruitment

We then investigated whether Gal3, ECAD and EPN3 form a functional complex at PM. MCF10A-EV control cells were incubated with recombinant His-tagged Gal3 at 4°C for 1 h to allow Gal3-ECAD binding at the cell surface. Gal3 pull-down assays performed on cell lysates confirmed the association of Gal3, ECAD, and EPN3 under basal conditions (**Fig. 3A**). In contrast, clathrin and AP2 were not enriched in the pull-down assays (**Fig. S6A**). Upon EPN3 overexpression, the binding of ECAD to His-Gal3 was markedly enhanced (**Fig. 3A, right**), consistent with the observed increase in Gal3-mediated ECAD endocytosis.

**Figure 3.**
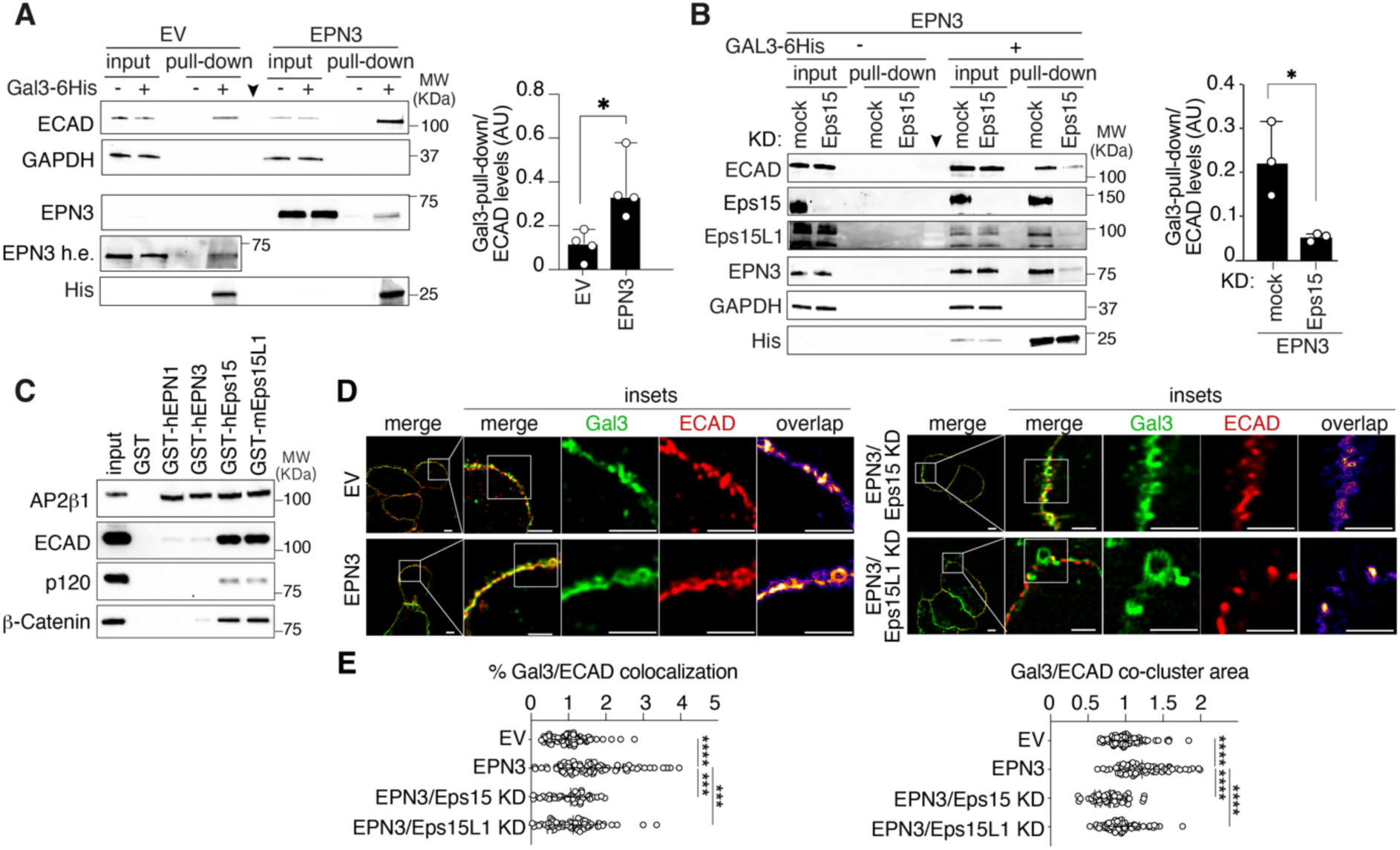
An EPN3-Eps15/L1 complex regulates Gal3-ECAD recruitment and clustering at the PM. **A)** Left, the transfectants indicated on top were incubated (+) or not (-) with recombinant human Gal3-6His, lysates were pulled-down with His-Fab trap and IB as shown on the left; The arrow marks the lane loaded with the molecular MW marker. Right, quantification of ECAD protein levels, normalized to His levels in the pull-down; n=4. **B)** Left, MCF10A-EPN3 cells were subjected to Eps15 KD (or mock) and then pull-down and IB as in (A). Right, quantification of ECAD protein levels as in (A); n=3. **C)** MCF10A lysates were pulled-down with the indicated human (h) or mouse (m) GST fusion proteins followed by IB as shown on the left. Input is 1/800^th^ of the pull-down fraction (except for AP2B1 which is 1/100^th^). **D)** Representative images of Gal3-ECAD clusters at the PM in MCF10A-EV, -EPN3, -EPN3/Eps15 KD and -EPN3/Eps15L1 KD cells by structured illumination microscopy (SIM); Gal3-488 (green), ECAD (red). Scale bars, 5 μm. **E)** Left, ECAD and Gal3 colocalization in the experiment shown in (D). Results are reported as a percentage of the total objects, normalized to EV. Right, dimension of Gal3/ECAD co-clusters, normalized to EV. N (fields of view): EV=80 (n=8), EPN3=78 (n=8), Eps15L1 KD=60 (n=6), Eps15 KD=40 (n=4). p-values are as in Fig. 1. In the various IB, GAPDH is used as negative control for IP and loading control for inputs and MWs are shown on the right in KDa.

Notably, both Eps15 and Eps15L1 were retrieved in the Gal3 pull-down alongside ECAD and EPN3 (**Fig. 3B**). The individual KD of either Eps15 or Eps15L1 in MCF10A-EPN3 cells markedly impaired the recruitment of ECAD and EPN3 to His-Gal3 (**Fig. 3B, right**, and **Fig. S6B**), indicating that Eps15 and Eps15L1 are critical for complex formation. Interestingly, the recruitment of one adaptor in the Gal3 pull-down was affected by depletion of the other, suggesting that Eps15 and Eps15L1 act as a molecular bridge between ECAD and EPN3, likely as heterodimers or higher-order oligomers, as previously proposed^27–29^. Supporting this model, the individual KD of either adaptor was sufficient to inhibit EPN3-induced ECAD endocytosis (**Fig. 1B**). Furthermore, GST pull-down assays using recombinant Eps15, Eps15L1, EPN3, or EPN1 as baits showed that ECAD binds directly to Eps15 and Eps15L1, but not to EPN3 or EPN1, whereas AP2 binding was equal across all conditions (**Fig. 3C**), reinforcing the idea that Eps15/L1 can bridge EPN3 to ECAD.

An unresolved question is how EPN3 overexpression facilitates the extracellular binding of Gal3 to ECAD through intracellular interactions with Eps15/Eps15L1. Using structured illumination microscopy (SIM), we observed that EPN3 overexpression significantly increased both the number and size of Gal3-ECAD clusters at the PM (**Fig. 3D, E**), an effect that was reversed by either Eps15 KD or Eps15L1 KD (**Fig. 3 D, E**). These findings suggest that EPN3, via its dual Eps15/Eps15L1 interaction surfaces, promotes ECAD clustering at the PM, thereby enhancing Gal3 recruitment and initiating ECAD internalization (**Fig. S6C**).

### EPN3 overexpression in the mouse mammary gland induces aberrant morphogenesis

To evaluate the pathophysiological relevance of EPN3-induced ECAD endocytosis, we generated keratin 5 (K5)-specific conditional EPN3-KI mice (**Fig. 4A**). These mice overexpress the human *EPN3* transgene in epithelial tissues, including the mammary epithelium (**Fig. 4B-C**). EPN3 expression in mammary epithelial cells derived from EPN3-KI mice was comparable to that observed in BT474 BC cells, which harbor *EPN3* gene amplification (**Fig. 4B**). Immunohistochemical (IHC) analysis confirmed *EPN3* transgene expression throughout the mammary duct epithelium, encompassing both the basal/myoepithelial (K14-positive) and luminal (K8-positive) layers (**Fig. 4C; Fig. S7A**).

**Figure 4.**
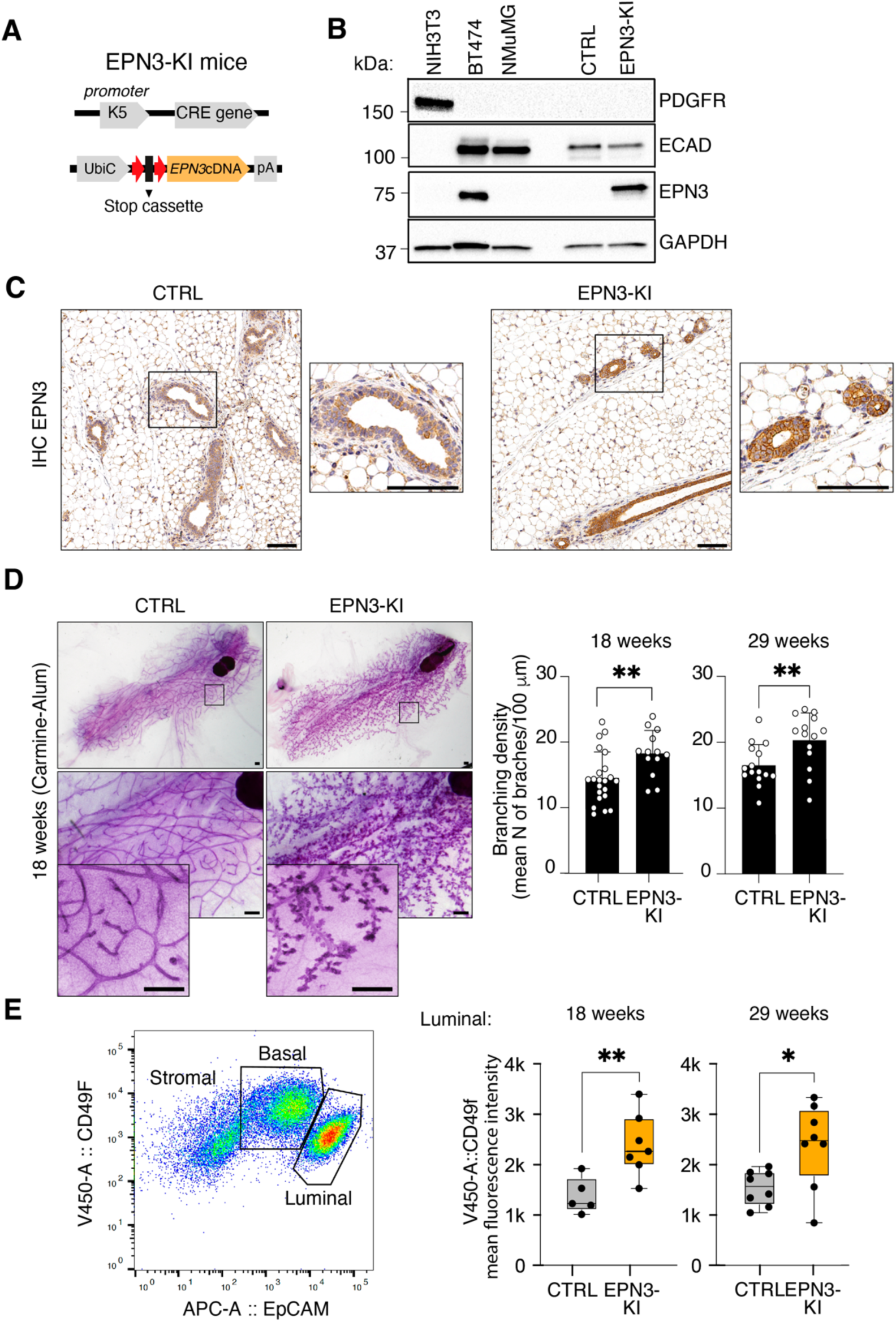
EPN3-KI mice exhibit aberrant mammary gland morphogenesis. **A)** Schematic of the breeding strategy used to generate EPN3-KI mice by crossing the FVB-Keratin 5 (K5)-CRE and FVB-EPN3-KI mice. K5 promoter-driven expression of CRE recombinase in epithelia induces site-specific recombination at the loxP sites (red arrows) resulting in removal of the stop cassette between the UbiC promoter and the human *EPN3* gene. **B)** IB analysis of EPN3 expression in pooled mammary epithelial cell populations isolated from two CTRL or two EPN3-KI mice, and in the indicated cell lines. GAPDH was used as loading control. MWs are indicated on the left. **C)** Representative IHC images of EPN3 expression in the mammary glands of CTRL *vs*. EPN3-KI mice. Boxed areas are magnified on the right. Bar, 100 μm. **D)** Left, representative images of whole-mount sections of mammary glands from 18-week-old CTRL and EPN3-KI mice; Bar, 500 μm. Right, quantification of mammary gland ductal branching density (mean number of branches per 0.1 mm section along the fat pad from the lymph node). N (animals); left, 18 weeks (CTRL=22; EPN3-KI=13); right, 29 weeks (CTRL=15; EPN3-KI=14). **E)** Mammary epithelial cells were isolated based on EpCAM expression. Left, representative bi-parametric FACS analysis of CD49f and EpCAM expression. The stromal, basal, and luminal subpopulations are indicated. Right, box plots showing the mean fluorescence intensity of the CD49f marker in luminal mammary epithelial cells in CTRL *vs*. EPN3-KI mice at 18 weeks (CTRL, N=5; EPN3-KI, N=7) and 29 weeks (N=8 per condition). p-values are as in Fig. 1.

EPN3-KI mice were born at Mendelian ratios and exhibited no overt developmental abnormalities or signs of mammary tumor formation or preneoplastic lesions up to 12 months of age. However, notable changes in mammary gland architecture and branching morphogenesis were observed. In virgin EPN3-KI females at late puberty (18 weeks) and adulthood (29 weeks), we detected mammary gland hyperplasia, increased secondary and tertiary ductal branching, and the formation of alveoli-like structures, compared to age-matched controls (**Fig. 4D; Fig. S7B**).

Given that enhanced ductal branching has been associated with an altered balance between myoepithelial/stem-like and luminal cell compartments^30, 31^, we analyzed these cell populations in the mammary glands from 18- and 29-week-old mice. Mammary epithelial cells were isolated based on EpCAM expression and immunophenotypically characterized by flow cytometry using combinations of EpCAM and integrin-α6 (Itga6)/CD49f (**Fig. 4E, left**)^32^. While no significant differences were observed in the overall proportions of luminal and basal populations (**Fig. S7C**), EPN3-KI luminal cells displayed a marked increase in Itga6/CD49f PM signal relative to controls (**Fig. 4E, right**). This result suggests that EPN3 overexpression may alter the expression or localization of markers involved in mammary epithelial cell differentiation, migration and/or adhesion. Notably, integrin-mediated interactions with the extracellular matrix have been shown to promote ductal branching during mammary gland development and lactation^33, 34^, implicating a potential mechanistic link between EPN3 activity and morphogenetic remodeling of the mammary epithelium.

### EPN3 overexpression induces upregulation of EMP and alveologenesis/lactation-like signatures

To gain molecular insights into the hyperbranching phenotype, we performed single-cell RNA sequencing (scRNA-seq) on viable EpCAM-positive cells isolated from pooled 18-week-old EPN3-KI and control mice (**Fig. 5A**). High-quality scRNA-seq data were obtained from 7,405 EPN3-KI cells and 6,783 control cells (**Fig. 5B**). UMAP clustering, based on 10 components, revealed 12 distinct clusters in both control and EPN3-KI cells, which were grouped into three principal epithelial categories (**Fig. 5C**). To classify these populations, we analyzed the expression of canonical mammary markers, as previously reported^35–37^, and identified them as basal/myoepithelial (BM), luminal progenitor (LP), and luminal hormone-sensing (LH) cell populations (**Fig. 5D, Fig. S8A-B** and **Table S2**). Consistent with FACS analysis (**Fig. S7C**), no significant differences in the distribution of cells across these populations were observed between control and EPN3-KI samples (**Fig. S8C**). However, we observed a selective upregulation of Itga6/CD49f expression in the LP population (**Fig. S8D**).

**Figure 5.**
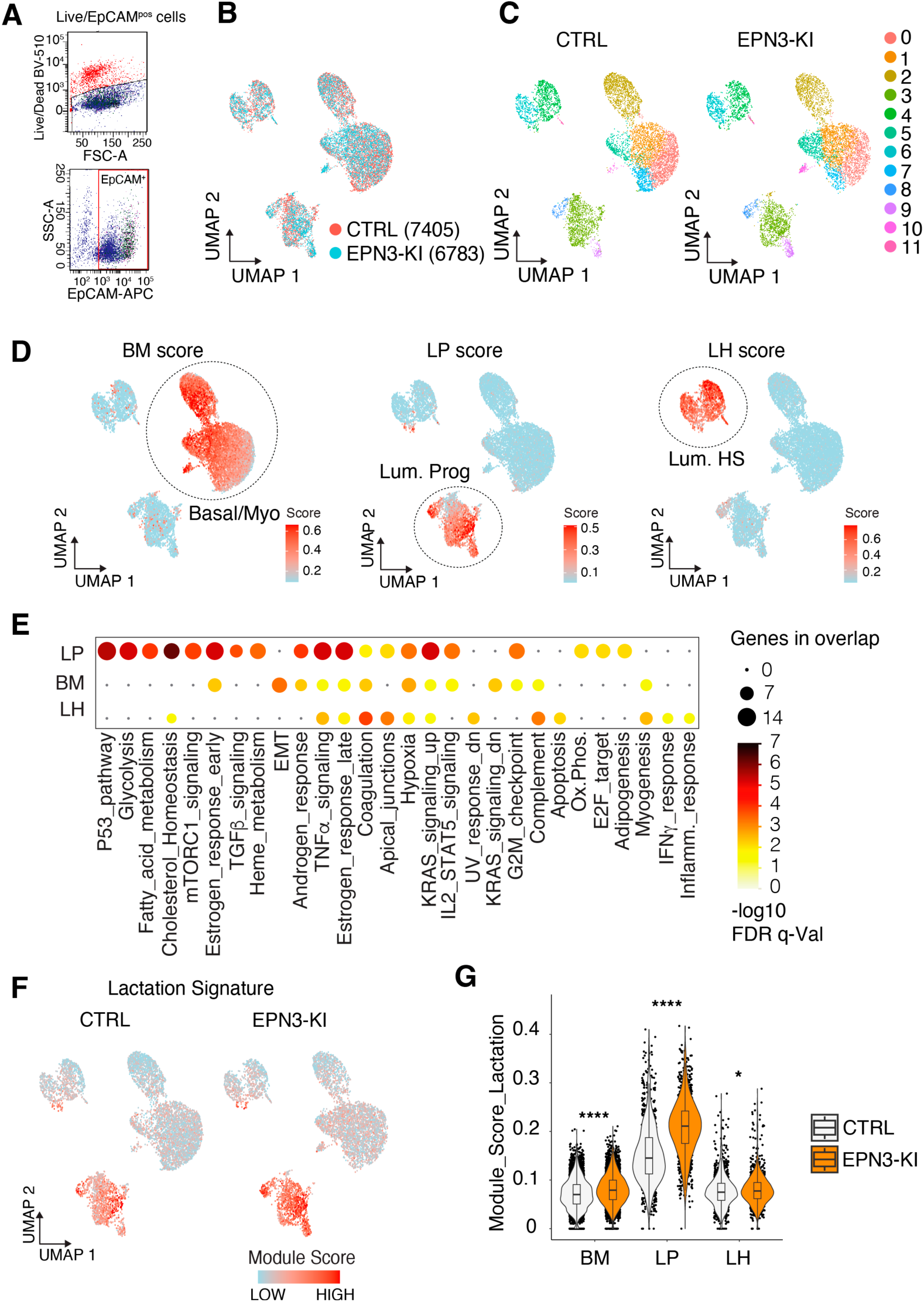
EPN3-KI mammary subpopulations exhibit upregulation of EMP and lactation/alveologenesis gene signatures. **A)** Mammary cells from a pool of 18-week-old CTRL (N=5) and EPN3-KI (N=7) mice were FACS-sorted to isolate viable EpCAM-positive cells for scRNA-seq analysis. Cells were stained with Live/Dead BV510 to exclude non-viable cells (top panel; BV510, Brilliant Violet 510 fluorochrome; FSC-A, forward scatter area). EpCAM expression was detected using allophycocyanin (APC)-conjugated antibody (bottom panel; SSC-A, side scatter area). Representative FACS plots show gating strategy. **B)** Unsupervised UMAP plot showing mammary epithelial cells populations: CTRL, red; EPN3-KI, blue. **C)** UMAP representation of CTRL and EPN3-KI cell clusters (0-11) identified by unbiased clustering. **D)** UMAPs showing the canonical expression level of basal/myoepithelial (BM), luminal progenitor (LP), luminal hormone-sensing (LH) gene signatures as “scores” (**Table S2**). **E)** Dot plot showing the enrichment of Hallmark pathways (mSigDb database, GSEA) in the list of significantly upregulated genes in EPN3-KI *vs*. CTRL subpopulations (BM, LP and LH). Circle size depicts the number of overlapping genes with a given pathway, and the color key denotes the -log_10_ FDR q-value overlap. **F)** Left, Expression of the lactation signature was measured in scRNA-seq profiles based on the levels of 30 markers ^60^ ^36^. Right, violin plot reports the distribution of the indicated module score by population; p-value was calculated with Wilcoxon rank sum test, comparing each cluster in the EPN3-KI *vs.* CTRL: **** <0.0001, * <0.05.

Differentially expressed genes (DEGs) within the BM, LP, and LH populations from EPN3-KI and control mice were identified. This analysis revealed a striking transcriptional upregulation, with 1,164 genes upregulated in the three subpopulations, compared to 195 genes downregulated (**Fig. S9A-C**; see also **Table S3**). Functional enrichment analysis of upregulated genes demonstrated notable transcriptional upregulation of EMT-related genes, including cell adhesion and extracellular matrix remodeling proteins (e.g., Ccn family proteins, metallo-proteases, collagens, and ECM modelers), specifically within the BM population (**Fig. 5E**, **Fig. S9D** and **Table S4**). Notably, canonical EMT transcription factors and markers, such as *Snai2*, *Mmp14*, *Vimentin*, and *Cd44* exhibited significant, albeit modest, upregulation in EPN3-KI BM cells compared to controls (**Fig. S9D**) and with respect to LP population. This expression pattern suggests activation of an EMP program in BM population, rather than a full EMT program, aligning with previous findings in EPN3-overexpressing MCF10A cells ^10^. Additionally, although ECAD expression was not downregulated at the transcriptional level (there was actually a slight, yet significant, increase, **Fig. S9E)**, there was a small yet reproducible decrease in ECAD protein levels (**Fig. 4B, Fig. S9F**), consistent with the establishment of EMP.

Functional enrichment analysis also revealed upregulation of signaling pathways (e.g., TGFβ and estrogen) and metabolic pathways (e.g., glycolysis, mTORC, cholesterol homeostasis, and fatty acid metabolism) specifically within the LP population (**Fig. 5E**). In addition, “lactation” represented one of the most highly upregulated pathways following EPN3 overexpression in LP cells as measured by module score, while a minor, albeit significant, upregulation was observed in BM cells (**Fig. 5F and Table S2**). These signatures were driven by the expression of genes typically associated with pregnancy and lactation, included caseins (*Csn1s1, Csn1s2a, Csn2, Csn3*), lactose synthase (*Lalba*) and milk proteins (*Wap)* (**Table S3**), all of which were among the top upregulated genes in LP in EPN3-KI mice **(Fig. S9A-C)** that had never undergone pregnancy.

Finally, module score analysis revealed that the BM, LP, and LH clusters contained subpopulations of cells with high proliferative scores, with EPN3-KI mice exhibiting approximately a three-fold increase in “cycling” cells compared to controls (**Fig. S10A**). This increase in proliferating cells induced by EPN3 overexpression was further confirmed by IHC analysis of Ki67-positive cells (**Fig. S10B**).

Taken together, these findings indicate that EPN3 overexpression promotes a morphogenetic phenotype resembling alveologenesis and lactation, characterized by enhanced proliferative activity and the activation of EMP programs. These alterations likely contribute to the promotion of branching and invasive behavior in mammary tubules within the mammary fat pad.

### Increased ECAD endocytosis drives EPN3-dependent phenotypes in primary mammary epithelial cells

A central mechanistic question is whether the observed *in vivo* phenotypes are directly associated with the role of EPN3 in driving ECAD endocytosis. To address this question, we performed ECAD endocytosis assays in live primary mammary epithelial cells isolated from control and EPN3-KI mice, in conjunction with Transwell invasion assays (**Fig. 6A**). We detected a significant increase in ECAD endocytosis in EPN3-KI cells *vs*. controls, which was inhibited by treatment with the competitive galectin inhibitor lactose or the specific Gal3 inhibitor I3 (**Fig. 6B**). This finding confirms a Gal3-driven ECAD internalization mechanism, as observed in MCF10A-EPN3 cells. Notably, this mechanism was also evident in control mammary epithelial cells, which exhibited sensitivity to the same inhibitors (**Fig. 6B**), supporting the notion that EPN3 overexpression amplifies a physiological mechanism rather than inducing a novel one. The increased ECAD endocytosis in EPN3-KI cells was accompanied by a corresponding increase in invasive capacity, which was reversed upon treatment with either lactose or I3 (**Fig. 6C**), thereby confirming that ECAD endocytosis contributes to the invasive phenotype.

**Figure 6.**
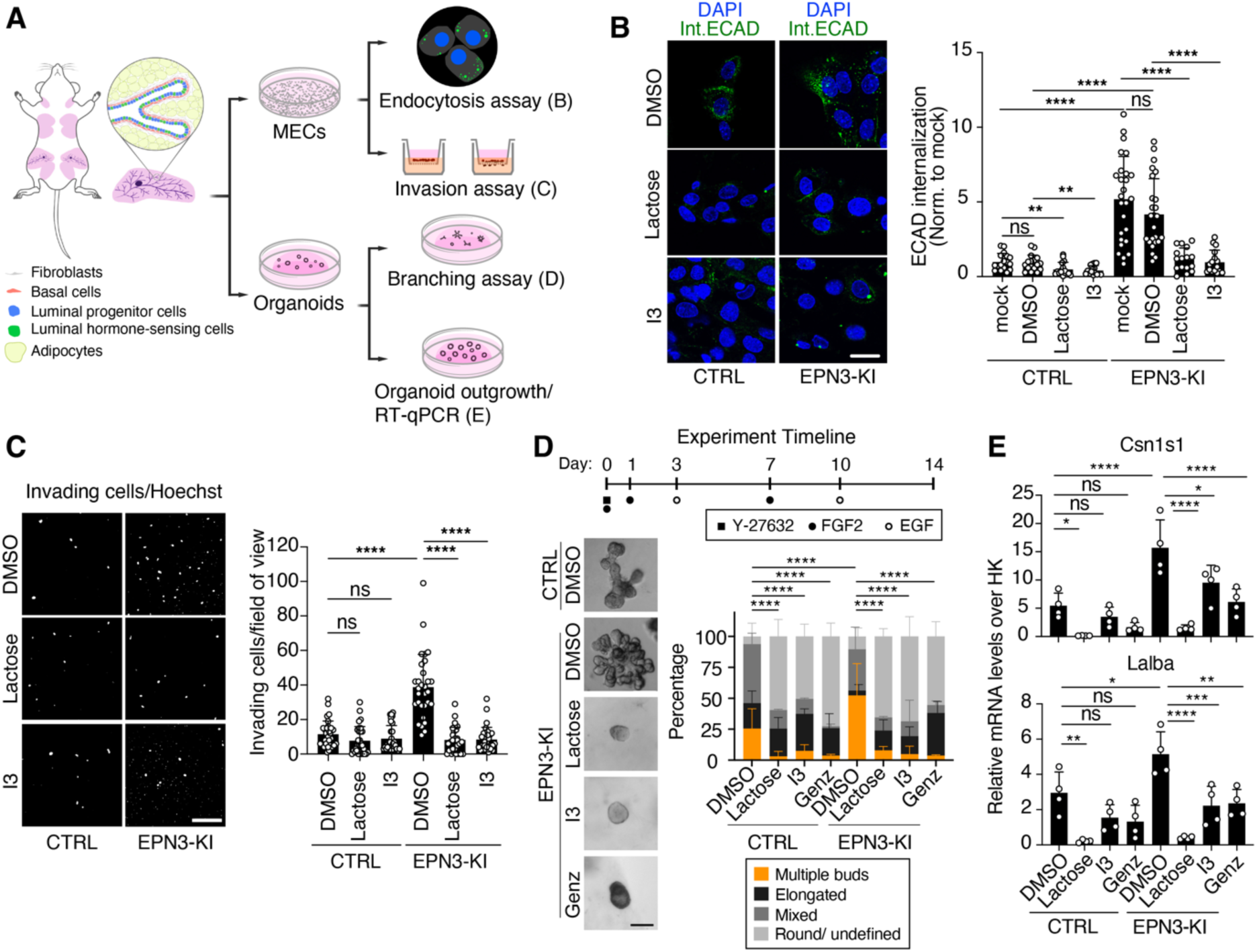
EPN3-mediated ECAD endocytosis drives invasive and alveologenesis phenotypes in primary mammary epithelial cells. **A)** Schematic summarizing the purification of murine primary mammary epithelial cells (MECs) and mammary organoids from 18-week-old mice used for endocytic, invasion, branching and organoid outgrowth assays. **B)** MECs were plated on collagenated µ-Slide 8 Well culture slides and pre-treated with (+)-Lactose (100 mM, 1 h) or I3 (20 µM, 10 min), before monitoring ECAD internalization *in vivo* by IF at 37°C for 30 min. Treatments were maintained during the internalization assay. Left, representative IF images showing internalized ECAD (green) and DAPI (blue) are shown. Bar, 20 µm. Right, quantification of relative ECAD fluorescence intensity/cell normalized to mock control. N (mice CTRL or KI) >10, n=3; p-values (Unpaired Student’s t-test, two-tailed): **** <0.0001; ** <0.01; ns, not significant. **C)** Transwell Matrigel invasion assay. Left, representative fields of view of invading cell nuclei stained with Hoechst; bar, 200 µm. Right, quantification of invading cells expressed as the number of invading cells per field of view. N (mice): CTRL or KI=8; N (fields of view)=30, n=3; p-values (Unpaired Student’s t-test, two-tailed): ****, <0.0001; ns, not significant. **D)** Mammary organoids were grown in Matrigel/collagen mixture and stimulated with the indicated factors following the timeline shown on top, in the presence or absence of (+)-Lactose (100 mM), I3 (20 µM), or Genz-123346 (4 µM). Left, representative images; bar, 100 µm. Right, stacked bar graph showing the morphological classification of organoids based on four categories: multiple buds, elongated, mixed, or round/undefined. CTRL: mock (n=4); Lactose (n=2); Gal3-I3 (n=2); Genz (n=2). EPN3-KI: mock (n=6); Lactose (n=2); Gal3-I3 (n=2); Genz (n=2). A contingency test was performed on the stacked bar graph, p-values: ****, <0.0001. **E)** RT-qPCR analysis of lactation gene expression (*Csn1s1* and *Lalba*) in mammary organoids grown in Matrigel in the presence or absence of Lactose (100 mM), Gal3-I3 (20 µM), or Genz (4 µM), n= 4. Data are shown as relative mRNA levels over 3 HK genes, p-values (calculated with one-way ANOVA, Šídák’s multiple comparisons test): ****, <0.0001; ***, <0.001; **, <0.01; *, <0.05; ns, not significant.

To further investigate whether the morphogenetic changes observed in the mammary glands of EPN3-KI mice are directly linked to ECAD endocytosis, we established an *ex vivo* branching morphogenesis assay (**Fig. 6A and 6D**, **top**)^38^. Organoids from control mice formed various structural types, including elongated, branched structures that resemble *in vivo* mammary ducts, and multilobulated structures reminiscent of ducts containing multiple alveoli (**Fig. 6D**). In contrast, organoids derived from EPN3-KI mice exhibited a marked shift toward multilobulated structures, mirroring the increased alveologenesis observed *in vivo* (**Fig. 6D**). Inhibition of ECAD endocytosis, using lactose, I3, or Genz, resulted in a dramatic increase in the percentage of rounded or underdeveloped organoids accompanied by a reduction in the number of multilobulated structures in both control and EPN3-KI organoids (**Fig. 6D**). These results suggest that ECAD endocytosis plays a role in branching morphogenesis and alveologenesis, even within a physiological context. Notably, RT-qPCR analysis of organoid-derived RNA confirmed that EPN3 overexpression induces upregulation of casein (Csn1s1) and lactalbumin (Lalba) expression, which was reversed upon inhibition of ECAD endocytosis (**Fig. 6E**), further supporting the link between EPN3, ECAD endocytosis, and alveolar differentiation.

In summary, our findings demonstrate that EPN3 overexpression in the mouse mammary gland drives an invasive, alveologenesis- and lactation-like phenotype, which is mechanistically linked to the promotion of GL-Lect driven ECAD endocytosis. Importantly, this endocytic mechanism is targetable, providing a potential therapeutic avenue to reverse invasive behavior.

### EPN3 overexpression accelerates ERBB2-driven breast tumorigenesis and lung metastasis

We hypothesized that the invasive, lactation-like EMP phenotype observed in EPN3-KI mice could cooperate with other oncogenes to predispose the mammary gland to tumorigenesis. To test this hypothesis, we crossed the EPN3-KI mice with tumorigenic MMTV-NeuN mice, which express the rat homologue of the human ERBB2 oncogene, NeuN, under the control of the mouse mammary tumor virus (MMTV) promoter (**Fig. 7A**). We confirmed that the resulting EPN3-KI:MMTV-NeuN mice co-overexpressed EPN3 and NeuN in mammary epithelial cells (**Fig. 7B**).

**Fig. 7.**
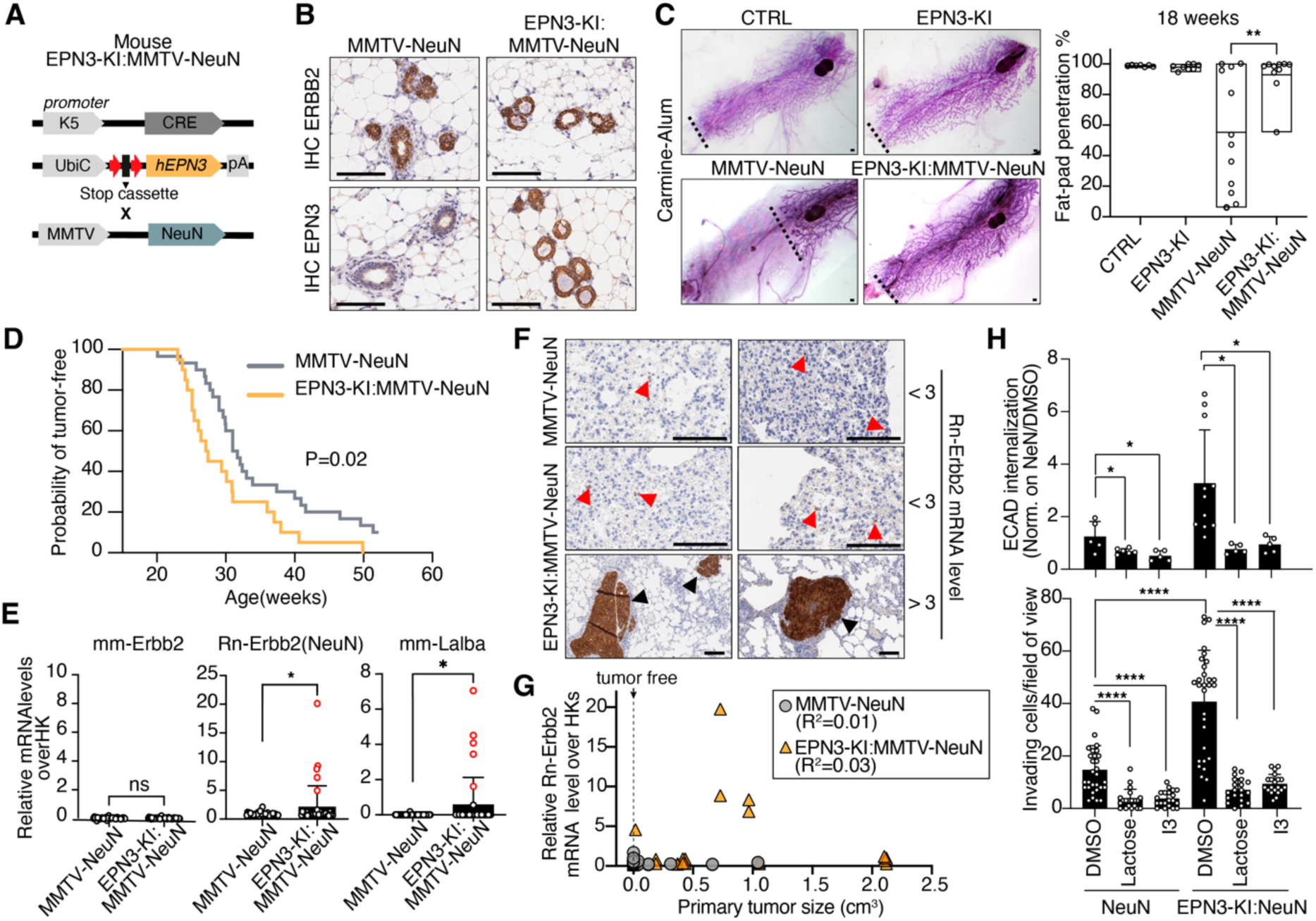
EPN3 overexpression accelerates ERBB2-driven breast tumorigenesis and lung metastasis. **A)** Schematic of the breeding strategy used to generate EPN3-KI:MMTV-NeuN mice. FVB.K5-Cre and FVB.EPN3-KI lines were first crossed to combine the Cre driver with the knock-in allele. Progeny carrying K5-Cre and EPN3-KI were then crossed with the MMTV-NeuN mice. **B)** Representative IHC images showing ERBB2 (NeuN) and EPN3 expression in mammary epithelial ducts of 18-week-old mice. Bar, 100 μm. **C)** Left, representative images of whole-mount sections of mammary glands from 18-week-old CTRL, EPN3-KI, MMTV-NeuN and EPN3-KI:MMTV-NeuN mice; Bar, 500 μm. Dotted lines indicate the limit of fat-pad penetration. Right, quantification of mammary fat-pad penetration (as distance from the lymph nodes) shown as percentage of the full-length of the fat pad (CTRL, N=7; EPN3-KI, N=8; MMTV-NeuN, N=12; EPN3-KI:MMTV-NeuN, N=10). **D)** Kaplan-Meier analysis of tumor-free survival in a cohort of EPN3-KI:MMTV-NeuN mice (N=20) and MMTV-NeuN mice (N=30), over a 1-year period. Time zero, when all mice were tumor-free, was set at 15 weeks. An event was recorded when tumors first became palpable. Survival curves were compared using the log-rank (Mantel-Cox) test, p=0.02. **E)** RT-qPCR analysis of endogenous Mm-Erbb2, transgenic Rn-Erbb2 (NeuN) and Mm-Lalba mRNA levels in lung tissues collected at 28–30 weeks of age from EPN3-KI:MMTV-NeuN (N=38) and MMTV-NeuN (N=44) mice. Results are expressed as relative to housekeeping (HK) genes (*Gapdh, Tbp, Gusb*). Samples with Rn-Erbb2 levels >3-fold higher than HK genes are shown in red. **F)** Representative anti-ERBB2 IHC staining of lung tissue samples used in (E). Upper panels: MMTV-NeuN lung samples with scattered metastatic cells (red arrowheads) and Rn-Erbb2 mRNA levels < 3. Middle panels: EPN3-KI:MMTV-NeuN lung samples with scattered metastatic cells (red arrowheads) and Rn-Erbb2 mRNA levels < 3. Lower panels: EPN3-KI:MMTV-NeuN lung samples with macro-metastatic lesions (brown areas) and Rn-Erbb2 mRNA levels >3; bar = 100 μm. **G)** Simple linear regression of the correlation between primary tumor size (in cm^3^) and relative Rn-Erbb2 levels over HK in lung tissue samples from MMTV-NeuN and EPN3-KI:MMTV-NeuN used in (E). Dotted line indicates the absence of palpable tumors (smallest tumor from EPN3-KI:MMTV-NeuN was 3x4 mm). **H)** Top, ECAD internalization was monitored by IF (37°C for 30 min) in primary MECs purified from NeuN (N=4) and EPN3-KI:MMTV-NeuN mice (N=4) and pre-treated with (+)-Lactose (100 mM, 1 h), I3 (20 µM, 10 min) or vehicle (DMSO). Quantification of ECAD fluorescence intensity per condition normalized to NeuN DMSO control. N (cells)>50 (n=2). Bottom, Transwell Matrigel invasion assay in the presence of Lactose (100 mM), I3 (20 µM) or vehicle (DMSO). Quantification of invading cells is expressed as the number of invading cells per field of view. N (mice): (NeuN, DMSO, N=8; EPN3-KI:MMTV-NeuN, DMSO, N=7; NeuN, I3/Lactose, N=6; EPN3-KI:MMTV-NeuN, I3/Lactose, N=6; N (fields of view): NeuN, DMSO=30 (n=3), I3=20 (n=2), Lactose=20 (n=2); EPN3-KI:MMTV-NeuN, DMSO=30 (n=3), I3=20 (n=2), Lactose=20 (n=2). p-values in all panels were calculated with Unpaired Student’s t-test, two-tailed: * p < 0.05; **, <0.01; ****, <0.000; ns, not significant.

The MMTV-NeuN mouse model was selected for several reasons. First, these mice develop spontaneous, monofocal mammary tumors with a low incidence of lung metastasis, thus providing a model that closely mirrors critical aspects of human BC progression^39, 40^. Moreover, our previous data suggest potential cooperation between EPN3 and ERBB2 in BC development: (i) EPN3 KD reduced the tumorigenicity of cells co-amplified for EPN3 and ERBB2 (e.g., BT474); (ii) EPN3 overexpression in the ERBB2-amplified HCC1569 BC cell line increased tumorigenicity in xenograft models and induced an EMP phenotype, characterized by disruption of ECAD-based adherens junctions; and (iii) approximately 50% of human BCs with EPN3 amplification also show co-amplification of ERBB2^10^.

While MMTV-NeuN mice exhibited a marked defect in mammary gland development, characterized by reduced mammary duct growth and impaired fat pad invasion (**Fig. 7C**), consistent with previous studies^41, 42^, we found that this phenotype was reversed in EPN3-KI:MMTV-NeuN mice (**Fig. 7C**). This effect is in line with our observations of upregulated migratory and invasive phenotypes in MCF10A-EPN3 cells and primary mammary epithelial cells (**Fig. 6C**)^41, 42^. In addition, EPN3-KI:MMTV-NeuN mice exhibited an earlier tumor onset compared to MMTV-NeuN control mice (**Fig. 7D**).

To assess the impact of EPN3 overexpression on metastasis, we developed an RT-qPCR assay that quantifies metastatic burden by detecting the relative levels of rat *NeuN* (Rn-Erbb2) mRNA in tissues. The assay was validated by correlating RT-qPCR measurements of Rn-Erbb2 mRNA with ERBB2 IHC staining in lung tissues containing varying metastatic loads (**Fig. S11A**). Increasing Rn-Erbb2 levels correlated with both the size and number of metastatic lesions detected by IHC (**Fig. S11B, left**). As a control, we measured murine *Erbb2* (Mm-Erbb2) mRNA in the same lung samples and observed consistently low levels of expression across all samples (**Fig. S11B, right**).

To evaluate the incidence of mammary metastases, formalin-fixed, FFPE lung tissue blocks were collected from 28-30-week-old mice. A subset of EPN3-KI:MMTV-NeuN mice exhibited significantly elevated pulmonary mRNA expression of *Rn-Erbb2* and *Lalba* compared with MMTV-NeuN controls, indicative of increased dissemination of NeuN/Lalba-expressing tumor cells and consistent with the presence of macrometastatic lesions (**Fig. 7E**). IHC analysis confirmed the presence of ERBB2-positive macrometastases in these lung samples, whereas only isolated ERBB2-positive cells were detected in *Rn-Erbb2* and *Lalba* low-expressing specimens (**Fig. 7F**). Notably, we found no correlation between primary tumor size and the appearance of lung macrometastatic lesions (**Fig. 7G**). These findings support the emerging evidence suggesting that metastatic dissemination may occur early during tumor evolution, driven by factors beyond tumor size^43–45^.

Finally, primary mammary epithelial cells derived from EPN3-KI:MMTV-NeuN mice exhibited upregulated ECAD endocytosis and increased invasiveness compared to cells from MMTV-NeuN or CTRL mice (**Fig. 7H** and **Fig. S11C-D**). Importantly, both phenotypes were suppressed by treatment with lactose or I3 confirming the role of the endocytic GL-Lect mechanism (**Fig. 7H).**

In summary, our findings indicate that while EPN3 overexpression has a modest impact on primary BC development, it significantly enhances mammary cell invasion and metastatic dissemination to the lungs.

## Discussion

Aberrant endocytosis has been implicated in various steps of tumorigenesis; however, direct *in vivo* evidence of its role in cancer progression remains limited. Here, we demonstrate that a form of clathrin-independent ECAD endocytosis – *i.e.,* the GL-Lect mechanism dependent on glycosphingolipids, Gal3 and the endocytic adaptors EPN3 and Eps15/Eps15L1 – drives aberrant mammary gland morphogenesis and increases susceptibility to metastatic progression in BC. Notably, this endocytic mechanism can be pharmacologically targeted using inhibitors that reduce the invasive capacity of mammary epithelial cells *ex vivo*, offering a new avenue for BC therapy.

Mechanistically, EPN3 defines a unique endocytic module distinct from its paralog EPN1. Through a dual interaction interface comprising a canonical NPF triad in its distal region and a noncanonical DPF motif in its central region, EPN3 binds to Eps15/Eps15L1 heterodimers. Our data argue that Eps15 and Eps15L1 act as heterodimers in this process. This molecular configuration promotes the formation of multivalent EPN3-Eps15/Eps15L1 complexes at the PM, where they orchestrate ECAD clustering and facilitate extracellular Gal3 recruitment.

Gal3 is a β-galactoside-binding lectin that is abundantly found in the extracellular space and biological fluids although it lacks conventional secretion signals^46, 47^. Extracellular Gal3, initially monomeric, has been shown to oligomerize up to tetramers and to multivalently bind glycosylated proteins and GSLs, driving the formation of membrane nanodomains and tubular endocytic pits from which clathrin-independent carriers emerge [REF^48–50^ and Shafaq-Zadah M. *et al.*, Nat Comm*, in press*]. These are morphologically similar to the ones described for clathrin-independent carrier-GPI-anchored protein-enriched early endosomal compartment endocytosis ^16, 51^. ECAD contains four N-glycosylation sites, one of which made of complex N-glycans that could interact with Gal3^52, 53^. Because each ECAD moiety binds only one Gal3 monomer, efficient GL-Lect driven endocytosis would require ECAD clustering at the PM, enabling Gal3 tetramers to bridge ECAD and glycosphingolipids. We propose that EPN3 coordinates this process from the cytosolic side, acting as a molecular organizer that couples endocytic adaptor networks to extracellular Gal3-driven nanodomains (**Fig. S6C**). This model establishes a direct structural and functional bridge between cytosolic endocytic scaffolds and extracellular lectin-driven domain organization, revealing a new paradigm for cross-compartmental coordination of endocytosis.

At the tissue level, EPN3 overexpression in the mammary epithelium amplifies a physiological GL-Lect driven endocytic mechanism, leading to the induction of an EMP program. These molecular effects manifest as enhanced ductal branching and alveoli-like structures in virgin glands. Indeed, the endocytic remodeling of adherens junctions, along with polarity loss and EMP activation, drives epithelial invasion and mammary tissue remodeling^54–56^. In addition, EPN3-KI mammary glands exhibit significant metabolic alterations, particularly increased lipid and cholesterol metabolism. These changes are accompanied by the activation of a transcriptional program reminiscent of alveologenesis and lactation, especially within LP cells – the alveolar progenitors. During normal alveologenesis and lactation, alveolar cells undergo extensive remodeling of intercellular junctions through enhanced intracellular trafficking, which supports membrane expansion and increased lipid metabolism^57, 58^. Similarly, in EPN3-KI glands, aberrant alveologenesis likely results from increased ECAD endocytosis, leading to disruption of cell-cell junctions and loss of epithelial polarity. Both processes require substantial PM remodeling, driving the heightened demand for lipids and cholesterol. Together, these phenotypes suggest that in BC, EPN3 hyperactivation hijacks a physiologic developmental endocytic program to drive morphogenetic plasticity. In agreement, endogenous EPN3 is upregulated in a human lactation signature^59^ suggesting a potential physiological role of EPN3 in tissue remodeling during lactation.

Functionally, the EPN3-Gal3 axis is required for both invasion and branching morphogenesis *ex vivo*. Pharmacological inhibition of Gal3 or glycosphingolipid synthesis reverses ECAD internalization, suppresses invasion, and restores epithelial architecture, underscoring the causal role of GL-Lect driven endocytosis in these processes. Moreover, when combined with *NeuN/ERBB2* oncogenic activation, EPN3 overexpression accelerates tumor onset and promotes lung metastasis, potentially linking dysregulated ECAD endocytosis to metastatic competence *in vivo*. These findings reveal that aberrant engagement of an endogenous endocytic route can potentiate oncogenic signaling and phenotypic evolution toward invasion and dissemination. EPN3-driven metastasis occurs independently of tumor size, indicating that metastatic potential emerges early in tumor evolution. Consistent with this, we previously found that endogenous EPN3 accumulates at the invasive front of mammary ductal carcinomas during the *in situ*-to-invasive transition in both xenografts and patient samples^10^, possibly implicating the GL-Lect mechanism as an early driver of metastatic progression.

In conclusion, our results establish EPN3 as a molecular organizer of GL-Lect driven ECAD endocytosis and highlight endocytic remodeling as a critical interface between cell adhesion, morphogenesis, and cancer progression. Given the accessibility of this endocytic mechanism to extracellular inhibition, targeting Gal3-ECAD interactions may provide a means to counteract EMP-driven invasion in breast cancer.

## Acknowledgments

We thank Rosalind Gunby for critically editing the manuscript. We thank Alberto Gobbi and the Mouse facility (Cogentech Società Benefit Srl, Milan) for mice handling. We thank IEO Technological Units, in particular: Molecular and Digital Pathology Unit, Imaging, Mass Spectrometry, Biochemistry and Structural Biology Units and Cell Culture Unit. We thank Simona Ronzoni and Flow Cytometry Unit for technical support. We thank Christian Wunder for scientific insights. We thank Elena Maspero and Simona Polo for providing Eps15 GST-tagged constructs.

## Author contributions

Conceptualization: SS, PPDF. Methodology: AF, ISL, AB, CT, MQ, SF, AR, FM, GB, LS, GC, AC, MSZ, ED, BG, GJ. Investigation: AF, ISL, AB, CT, MQ, SF, AR, GG, FM, INC, RC, GB, LS, GC, AC, BG, SP. Data analysis: AF, CT, AB, ISL, SK, SC, SF. Resources: SS, PPDF, LS, GC, HL, UJN, LJ, GS. Mouse experiments authorization: SS, CT, AF, AB, MQ, MGM. Funding acquisition: SS, PPDF, GS, LJ, SP. Supervision: SS, PPDF, GS, LJ, SP. Writing – original draft: SS; Writing – review & editing: SS, PPDF, CT, ISL, GS, LJ, HL, UJN, MGM.

## Data and materials availability

All data are available in the main text or the supplementary materials. Reagents and materials used in this study are either commercially available or can be obtained upon request.

Raw single-cell RNA-seq data is deposited in the GEO public repository. Accession numbers: GSM9326068, GSM9326069.

## Supplementary Materials

### Materials and Methods

#### Generation of genetic mouse models

The inducible EPN3-KI was generated as previously described as in^1^ and were maintained in the in-house animal facility. Briefly, the KI strategy is based on the targeted insertion of the human EPN3 cDNA (NM_017957.3) into the *ROSA26 locus* under the human Ubiquitin C promoter (UbiC)^2^, flanked by a loxP-STOP cassette between the promoter and the coding sequence. The STOP cassette can be removed using Cre recombinase leading to the inducible expression of the transgene. The EPN3-KI mice were back-crossed into FVB background to obtain FVB/EPN3-KI mice for experiments and then, crossed with the FVB/K5-Cre mouse model resulting in the EPN3-KI mouse which overexpresses *EPN3* in epithelial compartments including the mammary gland epithelium^3^.

MMTV-NeuN transgenic mice (FVB.N-Tg(MMTVneu)202Mul/J, JAX stock #002376) were purchased from The Jackson Laboratory^4, 5^. EPN3-KI:MMTV-NeuN mice were obtained by crossing MMTV-NeuN mice with EPN3-KI mice in the in-house animal facility. MMTV-NeuN and EPN3-KI:MMTV-NeuN tumorigenic mice were monitored 2-3 times per week for palpable tumors. Tumor dimensions were measured with Vernier calipers and tumor volume was calculated using the formula: volume = (length × width^2^)/2.

#### Mouse study approval

All mice were maintained in a controlled environment at 18-23°C, 40-60% humidity and with 12 h dark/12 h light cycles in a certified animal facility under the control of the institutional organism for animal welfare and ethical approach to animals in experimental procedures (Cogentech OPBA). All animal studies were conducted with the approval of the Italian Minister of Health (27/2015-PR, 550/2020-PR and 17/2023-PR) and were performed in accordance with Italian law (D.lgs. 26/2014), which enforces Directive 2010/63/EU of the European Parliament and of the Council of September 22, 2010, on the protection of animals used for scientific purposes.

#### Cell cultures

All human breast cell lines were from the American Type Culture Collection (ATCC), which were kindly provided by Dr. John F Marshall (Barts Cancer Institute, London, UK). All human cell lines were authenticated at each batch freezing by STR profiling (StemElite ID System, Promega) and tested for mycoplasma by PCR and biochemical assay (MycoAlert, Lonza).

MCF10A cells were cultured in a 1:1 mixture of DMEM and Ham’s F12 medium (Gibco, Life Technologies), supplemented with 5% horse serum (Invitrogen), 2 mM L-glutamine, 20 ng/ml human EGF (Invitrogen), 100 ng/ml cholera toxin, 10 μg/ml insulin and 500 ng/ml hydrocortisone (Merck Life Science). BT474 cells were cultured in DMEM supplemented with 10% FBS (South America origin) and 2 mM L-glutamine (Euroclone). NMuMG cells were cultured in DMEM supplemented with 10% FBS and stable glutamine, and 10 µg/mL insulin. NIH-3T3 cells were cultured in DMEM supplemented with 10% calf serum (CS, calf origin) and stable glutamine. HEK293T cells were cultured in DMEM supplemented with 10% FBS and stable glutamine. All cells were cultured at 37°C in a humidified atmosphere containing 5% CO_2_.

MCF10A cells infected with stably overexpressing pBABE-EPN3 retroviral construct or with pBABE empty vector (EV) as previously described as in^1^. Alternatively, MCF10A cells were infected with inducible pSLIK-neo EV or pSLIK-neo constructs overexpressing EPN1 or EPN3 full-length proteins or EPN1/EPN3 chimeras. Cells overexpressing chimeras or full-length proteins were selected with 150 μg ml^-1^ neomycin as previously described for MCF10A-pSLIK-neo EPN3 and -pSLIK-neo EV in^1^. To induce expression of proteins, cells were treated with a range of 100 to 600 ng ml^-1^ doxycycline for 24 h. Reagents used for tissue culture are listed in Table S5.

#### Isolation of primary mammary epithelial cells (MECs)

To isolate primary MECs, mice were sacrificed and inguinal and thoracic mammary glands were excised and subjected to overnight mild enzymatic digestion in medium DMEM-Ham’s F12 medium (Gibco, Life Technologies), 20 µg/ml Liberase™ (Roche), 150 U/ml Collagenase Type 3 (Worthington), 1 µg/ml human insulin, 1 µg/ml hydrocortisone (Merck Life science), 10 ng/ml hEGF (Invitrogen), 10 mM HEPES, and 1% Pen/Strep. Samples were incubated at 37°C in a humidified atmosphere containing 5% CO_2_.

Cell suspension was resuspended in PBS and centrifuged at 335 g for 5 min at room temperature (RT). A three-layer suspension was formed, with the upper layer containing epithelial cells trapped in the fatty layer, a middle layer containing stromal cells that was removed by aspiration and a pellet containing epithelial organoids/MECs and red blood cells. The upper fatty layer and the pellet were harvested and incubated at 37°C with 0.25% Trypsin-EDTA solution for 20 min, pipetting the suspension every 10 min. Trypsin activity was then inhibited with 1X trypsin inhibitor at 1:1 ratio (Merck Life science). If necessary, the suspension was incubated at 37°C with 10 U/ml DNAse I (Merck Life science) for 2 min at RT followed by centrifugation at 335 g for 5 min. The supernatant was removed, and red blood cells were removed with ACK Lysing buffer (Gibco). Samples were then centrifuged at 335 g for 5 min to obtain a pellet of purified MECs. Cells were immediately used for further analyses (see FACS staining and scRNA-seq) or plated with Mammary Epithelial Cell Growth Medium (MEGM, Lonza) on collagen I-coated plates (Corning® BioCoat™) for 24 h cells at 37°C in a humidified atmosphere containing 5% CO_2_ and then used for subsequent analyses (see ECAD endocytosis and Transwell invasion assay). A list of reagents used is provided in Table S5.

#### Endocytic inhibitors

The following inhibitors of endocytic players were used in this study: CK666 (Arp2/3 complex inhibitor, 50 µM, SML0006 Merck), EIPA (NHE inhibitor, 50 µM; used as macropinocytosis inhibitor, A3085 Merck), I3 (non-permeable Galectin-3 inhibitor, 20 µM, GalectoBiotech), Genz-123346 (ceramide synthase inhibitor, 4 µM, 5.38285 Merck), and (+)-Lactose (pan-galectin inhibitor, 100 mM, L3750 Merck). The 100 mM Lactose medium was prepared as follows: complete medium was diluted 1:1 with ddH₂O. Lactose or Sucrose (as control) was added and stirred. NaCl was added to a final concentration of 25 mM (50 mOsm) to maintain the correct osmolarity of the solution (300 mOsm).

For determining mouse galectin-3 affinity for I3, competitive fluorescence polarization experiments were performed with a PheraStarFS plate reader and PHERAstar Mars version 2.10 R3 software (BMG, Offenburg, Germany). Fluorescence anisotropy of the fluorescein-tagged 3,3’-dideoxy-3-[4-(fluorescein-5-yl-carbonylaminomethyl)-1H-1,2,3-triazol-1-yl]-3’-(3,5-di methoxybenzamido)-1,1’-sulfanediyl-di-b-D-galactopyranoside (0.02 µM) was measured in PBS as previously described^6, 7^ at 20°C with excitation at 485 nm and emission at 520 nm in the presence of mouse galectin-3 (0.50 µM) and different concentrations in duplicate of I3 (0.02-5.6 µM). Average Kd values and SEMs were calculated from nine single-point measurements showing between 20–80 % inhibition to give a Kd of 120±21 nM.

#### RNA interference (RNAi)

RNAi was performed with Lipofectamine RNAimax reagent from Invitrogen, according to manufacturer’s instructions. Cells were subjected to 2 cycles of transfection (reverse at day 0 and forward at day 1) with 8 nM of siRNA, with the following sequences:

AP2µ (Invitrogen) 5’-CAUUGACCCGAAAGGCAUCCACUG-3’

Eps15L1 (Invitrogen) 5’-CCAGAAGGUAAAGGGUUCUUGGACA-3’

EPS15 sipool (Biotech) gene ID 2060

Caveolin1 (Invitrogen) pool;

5’-AUUAUGAAGUGAAUGGCAGCCCCC-3’

5’-ACAUAACAAAUAUUCAGGCCCCC-3’

5’-UACUUGUACACUCCAUGGCCCCC-3’

5’-AGUUCUUCAUAUUUCUCUCCCCC-3’

Caveolin2 (Riboxx) pool;

5’-AUUCAUUUGAUUUGGUCCCCC-3’

5’-UUUAUAUUCACCAGUCCUGCCCCC-3’

5’-ACUUACUGUAUAAUUCUGGCCCCC-3’

5’-UACAUAAUCAGAACAGUGCCCCC-3’

RTN3 (Invitrogen) 5’-CCCUGAAACUCAUUAUUCGUCUCUU-3’

Gal3 sipool (Biotech) gene ID 3958

As a negative siRNA control, mock-silenced cells were used (CTR).

#### Generation of chimeric EPN3/EPN1 constructs

Chimeras of human EPN3/EPN1 proteins were generated by swapping EPN1 and EPN3 regions by PCR using pEN_Tmcs EPN3 previously described in ^1^ and pEN_Tmcs EPN1 plasmid which was generated by cloning EPN1 isoform C human cDNA from pBABE vector^1^ into pEN_Tmcs vector. In particular, each chimera was generated by performing two PCRs: a PCR using 5’ phosphorylated primers to amplify the desired insert region of one protein (*e.g* EPN3-ENTH, - ENTH-PR, -CR-DR, -CR, -DR) and a PCR using primers to amplify the rest of the desired regions (*e.g.* EPN1 regions) plus the entire backbone. The two PCR products for each chimera were run on a 0.8% agarose gel, and the bands were extracted according to the manufacturer’s instructions, using the kit Nucleospin gel and PCR-clean up (Macherey Nagel). DNA was quantified and ligation was performed using the Quick Ligation Protocol (Biolabs) with a ratio insert to vector 3:1. TOP10 bacteria were transformed, mini- and maxi-preps were performed using Nucleospin Plasmid Protocol or Nucleobond Xtra maxi (Macherey Nagel), screened and DNA was sequenced. Finally, recombination of pEN_Tmcs chimeras into pSLIK vector was performed using LR Clonase Gateway System (Thermo Fisher), STBL3 bacteria were transformed, maxi-preps were performed using Nucleobond Xtra maxi (Macherey Nagel) and DNA was sequenced.

A list of the primers used for the generation of each chimera is provided in Table S5.

#### IF internalization assays in MCF10A cells

MCF10A-EV and -EPN3 cells, were mock-treated or subjected to KDs of various endocytic players for 4 days. Cells were washed with PBS and EGF-starved for 3 h. Then, cells were incubated either with anti-ECAD Antibody (HECD1 at 2 µg/ml) for 1 h at 4°C or anti-CD44 (Hermes3 at 1 µg/ml) for 1 h at 4°C in EGF-starved medium. Cells were then shifted to 37°C in complete medium for the indicated time points. At each time point, cells were washed once with cold PBS and treated with acid wash solution (AW, glycine 0.1M in ddH_2_O, pH=2.2) on ice for 3 cycles of 45 sec to remove surface-bound ECAD or CD44 antibodies. Cells were fixed for 10 min with PFA 4% at RT, permeabilized with Triton X100 0.1%, followed by incubation with secondary Antibody (Alexa-488, Thermofisher 5 µg/ml, see Table S5) for 30 min to visualize internalized ECAD/CD44, and DAPI counterstained. In the case of treatments with endocytosis inhibitors, after EGF starvation, cells were either treated with vehicle control (DMSO) or pre-treated with the following inhibitors at 37°C: 20 μM I3 for 10 min, 50µM EIPA for 30min or 50 μM CK666 for 1 h. ECAD/CD44 endocytosis was followed as above. Inhibitors were present during all incubation steps.

For lactose treatment, cells were incubated with 100 mM (+)-Lactose for 1 h. As a control (mock), cells were treated with 100 mM sucrose, as previously described (see e.g.,REF^8^). Since mock (sucrose) and DMSO controls showed comparable results, only DMSO controls are displayed in some figures.

Cells were incubated with Alexa 488/647-Transferrin (Thermofisher, see Table S5 for specifications) and Alexa488-Shiga-toxin ligands at 37°C for 30 min. At each time point, cells were washed once with cold PBS and treated with AW (glycine 0.1M, pH=2.2) on ice for 3 cycles of 45 sec. Cells were fixed for 10 min with PFA 4% at RT and DAPI stained.

Images were acquired at confocal microscope (SP8, Leica or Nikon Ti2, equipped with Spinning Disk module) and processed with Image J software. Different fields of view for each sample were imaged and quantified by Integrated Density (IntDen) to evaluate internalized signal intensity which was divided by the number of cells per field of view.

#### ECAD internalization assays by FACS in MCF10A

ECAD internalization was performed as previously described in^1^. Briefly, MCF10A cells overexpressing EPN3 and EPN1 full-length proteins or EPN3-EPN1 chimeras were seeded and induced with doxycyline 200 ng ml^-1^ 24 h prior to the assay. Cells were then EGF starved for 3 h, incubated with an anti-ECAD antibody (HECD1, 2 µg/ml) for 1 h at 4 °C, followed by an anti-mouse AlexaFluor488-conjugated secondary antibody for 30 min at 4 °C. After a wash in medium, cells were incubated at 37°C in complete medium for the desired time points, with exception of time 0 samples, which were immediately fixed to evaluate the total amount of ECAD at the PM.

At the indicated time points, cells were washed once with cold PBS and treated with AW solution (0.1 M glycine, pH 2.2) on ice, three times for 45 seconds each. As a control for the efficiency of the acid wash, a time 0 sample treated with acid wash was collected. Cells were washed twice with PBS and detached using 0.25% trypsin for 15–20 min at 37°C. All cells were recovered in medium in a 15 ml Falcon, washed once with PBS, re-suspended in 500 µl PBS and fixed by adding 500 µl of formaldehyde 2% for 15 min on ice, and re-suspended in PBS with EDTA 2 mM before analysis with the FACS Celesta (BD).

To evaluate the amount of internalized ECAD, the mean fluorescence intensity of internalized ECAD for each time point was divided by the total ECAD fluorescence intensity. The time 0 sample treated with acid wash was used as the background. Samples stained with only the secondary antibody served as negative controls. Data analysis was carried out using FlowJo software (version 10.4.2, LLC). Briefly, doublets and cell aggregates were excluded based on forward scatter area (FSC-A) versus height (FSC-H). Cells with appropriate morphology were then gated using side scatter area (SSC-A) versus FSC-A.

#### ECAD internalization assays by IF in MECs

MECs were plated on collagen I coated 8-well slides (µ-Slide ibidi®), 1 day before the assay at 60-80% confluence. Cells were pre-treated with inhibitors: 100 mM (+)-Lactose (in MEGM) for 1 h at 37°C, 20 μM I3 for 10 min at 37°C or DMSO. Cells were incubated with the ECAD antibody (DECMA-1 at 5 µg/mL) for 1 h at 4°C and then ECAD internalization was followed at 37°C in presence or not of inhibitors/DMSO for 30 min. After this time point, cells were treated with AW solution (Glycine 0.1M, pH=2.2) for 3 cycles of 45 seconds. Cells were washed twice with PBS and fixed in 4% PFA at RT, followed by incubation with blocking buffer (5% Donkey Serum, 2% BSA, in 1X PBS). Incubation with Alexa-488/647-conjugated secondary antibody (Thermofisher, see Table S5 for specifications) was performed in blocking buffer and DAPI was used to counterstain nuclei. Cells were then analyzed by confocal microscope SP8 CSU (Leica). Images were acquired at confocal microscope (SP8, Leica or Nikon Ti2, equipped with Spinning Disk module) and processed with Image J software. Different fields of view for each sample were imaged and quantified by Integrated Density (IntDen) to evaluate internalized signal intensity which was divided by the number of cells per field of view.

#### Aspect ratio evaluation

MCF10A cells overexpressing EPN3 and EPN1 full-length proteins or EPN3-EPN1 chimeras were seeded on gelatin 0.1% coated coverslips and induced with doxycycline 24 h prior to the assay. Coverslips were then fixed for 8 min with 4% PFA and cells were stained with anti-phalloidin Rhodamine-conjugated (ThermoFisher, see Table S5) and nuclei were stained with DAPI. Images were acquired using SP8 Confocal Microscope 63X oil and used to evaluate aspect ratio of cells, *i.e.* the ratio between the long axis and short axis of each cell. A threshold of aspect ratio > 2.5 was considered.

#### FLAG co-immunoprecipitation

MCF-10A cells overexpressing EPN3 and EPN1 full-length proteins or EPN3-EPN1 chimeras were seeded and induced with a range of 100 to 600 ng ml^-1^ doxycycline 24 h prior to the assay. Cell lysates were then prepared in JS buffer (50mM Hepes pH 7.5, 150mM NaCl, 1% Triton X100, 5mM EGTA, 1.5mM MgCl2, 10% glycerol) supplemented with a protease inhibitor cocktail (Calbiochem) and cleared at 16000 g for 20 min. Protein supernatants were quantified and 1 mg of lysate was incubated with 6 μl of slurry anti-FLAG M2-conjugated beads (Merck) for 2 h at 4°C in a rotating wheel. Following centrifugation, beads were washed 4 times in JS buffer and proteins were eluted in 2X Laemmli buffer for 10 min at 95°C. Proteins were separated by SDS-PAGE and Coomassie stained to quantify the bait; accordingly, proteins were then prepared for SDS-PAGE followed by IB to detect co-immunoprecipitated proteins. Samples were run on separate gels to detect proteins with similar molecular weights. Gels were then sectioned into regions of interest to enable probing with different antibodies (see Table S5).

#### Immunoblot analysis

Cells and MECs were lysed by adding RIPA buffer (50 mM Tris–HCl, 150 mM NaCl, 1 mM EDTA, 1% Triton X-100, 1% sodium deoxycholate, 0.1% SDS), plus protease inhibitor cocktail (Calbiochem) and phosphatase inhibitors (20 mM sodium pyrophosphate pH 7.5, 50 mM NaF, 2 mM PMSF, 10 mM Na_3_VO_4_ pH 7.5). Lysates were clarified by centrifugation at 16,000 g for 20 min at 4°C. Protein concentration was measured by Bradford Assay (Biorad) and 5–50 μg of protein were run on 4-20% gradient pre-cast Gels (Biorad) and transferred using Trans-Blot (Biorad) according to manufacturer’s instructions. Filters were blocked with 5% milk diluted in TBS - 0.1% Tween20 (TBS-T) and then incubated overnight with primary antibody (see Table S5). Following 3 washes with TBS-T, filters were then incubated with the appropriate secondary antibody conjugated with horseradish peroxidase. After 3 more washes, the signal was detected with Chemidoc (Biorad) or iBright (Invitrogen) using ECL Western Blotting Detecting Reagents (Amersham), Clarity (Biorad) or SuperSignal West Femto (Thermo Fisher Scientific).

#### AlphaFold3 modeling

AlphaFold3 server (https://alphafoldserver.com)^9^ was used to predict the complexes between EPN1-DR (UniProt Q9Y6I3-3, residues 467–550) or EPN3-DR (UniProt Q9H201, residues 517–632) and EPS15 (UniProt P42566, residues 1–313) or EPS15L1 (UniProt Q9UBC2, residues 1–364). For each model, ten independent runs starting from different seeds were performed, each generating five models for a total of 50 models per complex. The percentage of models displaying two or three binding sites was calculated from these 50 predictions. All the predictions were computed using default AlphaFold3 settings. Molecular representations were prepared in PyMOL (Schrödinger, LLC. PyMOL Molecular Graphics System, Version 2.6.2). Predicted Aligned Error (PAE) plots of the represented models were generated using PAE viewer (https://pae-viewer.uni-goettingen.de/)^10^ and exported for representation.

#### Electron microscopy

##### Pre-embedding immunolabeling

Cells were plated on Mattek 35-mm glass bottom dishes at 70-80% of confluency, incubated with anti-ECAD (HECD-1) antibody, followed by incubation with rabbit anti-mouse, and, finally, with Protein-A Gold 10 nm (30 min incubation on ice/each step). Cells were then incubated at 37°C for 5 min, as indicated. As a control, one sample was kept at 4°C for 5 min to assess antibody binding. Cells were then washed in PBS and fixed for 1 h at RT in formaldehyde/glutaraldehyde 2.5%/1.2% in 0.1 M sodium cacodylate buffer pH 7.4 containing 0.5 mg/ml of ruthenium red. After quick washes with 150 mM sodium cacodylate buffer, the samples were post-fixed in 1.3% osmium tetroxide in a 33 mM sodium cacodylate buffer containing 0.5 mg/ml ruthenium red for 2 h at RT. Cells were then rinsed with 150 mM sodium cacodylate, washed with distilled water and stained with 0.5% uranyl acetate in dH20 overnight at 4°C in the dark. Finally, samples were rinsed in ddH_2_O, dehydrated with increasing concentrations of ethanol, embedded in Epon and cured in an oven at 60°C for 48 h. Ultrathin sections (70–90 nm) were obtained using an ultramicrotome (UC7, Leica microsystem, Vienna, Austria), collected, stained with uranyl acetate and Sato’s lead solutions, and observed in a Transmission Electron Microscope Talos L120C (FEI, Thermo Fisher Scientific) operating at 120 kV. Images were acquired with a Ceta CCD camera (FEI, Thermo Fisher Scientific).

##### Morphometry of NCE TIs and of CCPs

Morphometry was performed as previously described in ^11, 12^. Briefly, cellular profiles of thin sections of cells immunolabeled with anti-ECAD antibody (HECD-1) and stained with ruthenium red were acquired. For the quantification of the number of CCPs or TIs, gold particle clusters present in PM connected (ruthenium red-positive) structures of randomly selected cells were acquired at a nominal microscope magnification of ×22,000 (pixel size 0.6 nm). Gold clusters were assigned to one of the two categories based on the presence of a clathrin coat and the distance between PM and the tip of the invagination was measured using ImageJ. The number of ruthenium red-positive endocytic structures identified was divided by the PM length measured with ImageJ on acquired low magnification micrographs (nominal microscope magnification of ×1200) and expressed as a percentage of structures detected on PM.

#### Gal3 cell surface co-immunoprecipitation

Gal3 co-immunoprecipitation was performed as previously described^8^ with some modifications. Briefly, cells were seeded at 60% confluency and incubated for 10 min at 4°C in (+)-Lactose buffer (20 mM HEPES, pH 7.3; 200 mM (+)-Lactose; 45 mM NaCl; 5 mM KCl; 1 mM MgCl₂; and 1 mM CaCl₂) to remove endogenous Gal3 bound to the cell surface. Cells were then washed three times with PBS and incubated in serum-free medium, either alone or supplemented with 5 µg/ml Gal3–His, for 45 min at 4 °C. After incubation, cells were rinsed with PBS and lysed in TNE buffer (10 mM Tris-HCl, 150 mM NaCl, 0.5% NP-40 substitute, 0.5 mM EDTA, 1 mM PMSF) supplemented with protease inhibitor cocktail (Calbiochem). Lysates were kept on ice for 30 min by pipetting every 10 min and cleared by centrifugation at 17,000 g for 10 min at 4°C. The resulting supernatants were incubated overnight at 4°C with His Fab-Trap® (Proteintech), followed by four washes with lysis buffer. Proteins were eluted using 2× SDS sample buffer.

#### Gal3 co-binding and co-uptake assays

Gal3 co-uptake and co-binding assays were performed as previously described^13^.

##### Co-binding assay

Cells were seeded onto coverslips at 60% confluency and then incubated at 4°C for 30 min with 200 nM Gal3-Alexa488. After incubation, cells were washed with ice-cold PBS and subsequently incubated with 2 µg/mL of anti-ECAD antibody (HECD1) for an additional 30 min at 4 °C. Following antibody incubation, cells were fixed in 4% PFA prepared in PIPES buffer.

##### Co-uptake assay

Cells were subjected to sequential incubation with Gal3–Alexa488 and anti-ECAD as described in the co-binding assay. Then, cells were shifted at 37°C to allow internalization for 10 min. Cells were then washed three times for 5 min at 4°C with iso-osmolar lactose DMEM/F12 medium (150 mM lactose), followed by treatment with acid wash solution (glycine 0.1 M, pH 2.2) for 3 cycles of 45 seconds. Cells were then washed thoroughly with PBS and fixed in 4% PFA in PIPES buffer.

##### Immunostaining and mounting

Fixed cells were blocked with 2% BSA in PBS. For co-binding assays, secondary antibody staining with Alexa Fluor-conjugated anti-mouse IgG was performed in blocking buffer without cell permeabilization. For co-uptake or single-uptake assays, cells were permeabilized with 0.1% Triton X-100 in PBS, followed by blocking and secondary antibody staining as above. Nuclei were counterstained with DAPI for 5 min and coverslips were mounted using a glycerol-based mounting medium.

#### SIM (structured illumination microscopy)

Images were acquired using a CrestOptics DeepSIM (Nikon) equipped with an oil-immersion objective lens 60X (1.40 N.A.) and Yokogawa Spinning Disk Field Scanning Confocal System. The analyses were performed through ImageJ software; the percentage of colocalization was calculated using an object-based analysis [JACOP plugin^14^]. The same threshold used to calculate the objects for colocalization analysis, was used to determine the co-cluster area, defined as the area where the signal of ECAD and the signal of Gal3 overlaps.

#### Recombinant protein expression and purification

GST-fusion proteins were expressed in BL21 Rosetta *E. coli* cells by inducing expression for 6-8 h at 20°C with 0.3 mM IPTG. Cells were lysed in a buffer containing 100 mM Tris-HCl pH 7.6, 300 mM NaCl, 10% glycerol, 0.5 mM EDTA, 1 mM DTT, 0.2 mg/ml lysozyme, and a protease inhibitor cocktail (Complete, EDTA-free, Roche). Following lysis, samples were sonicated and the lysate was clarified by centrifugation at 20,000 g for 30 min. The resulting supernatant was incubated with Glutathione Sepharose 4 Fast-Flow beads (GE Healthcare) to capture the fusion proteins. After incubation, the beads were sequentially washed: i) 100 mM Tris-HCl pH 7.6, 300 mM NaCl, 0.5 mM EDTA, and 1 mM DTT; ii) the same buffer containing 1 M NaCl; and iii) the original 300 mM NaCl buffer. The beads were then resuspended in JS buffer for the pull-down assay.

Production of recombinant Gal3-6His was performed as previously described^8^, with some modifications. A plasmid encoding a C-terminal His-tagged version of WT Gal3 (pHisParallel2) was transformed into *E. coli* BL21 Rosetta cells and incubated overnight at 37°C. A single colony was inoculated in 50 ml of LB overnight at 37°C and subsequently inoculated in 1 L of LB at 37°C with shaking until OD₆₀₀ reached 0.6. Protein expression was then induced at 20°C overnight by adding IPTG to a final concentration of 600 µM. Following expression, all subsequent steps were performed at 4°C or on ice. Cells were harvested by centrifugation at 4,500 g for 20 min and resuspended in PBS containing 10 mM imidazole, 10% glycerol, DNAaseI (PanReac Applichem), EDTA-free protease inhibitor cocktail (Calbiochem) and 1 mM PMSF (Merck). Cells were lysed by sonication and the lysate was clarified by centrifugation at 70,000 g for 60 min. The cleared supernatant was applied to a HiTrap Chelating 5 ml column (Cytiva) pre-loaded with cobalt chloride, using an AKTA chromatography system (Cytiva). After a wash with lysis buffer supplemented with 10 mM (+)-Lactose, Gal3–His was eluted from the resin with PBS containing 500 mM imidazole. The eluate was concentrated with 10 kDa cut-off Amicon ultra centrifugal filters (Millipore) and further purified by size-exclusion chromatography on a Superdex 75 16/60 column equilibrated in PBS. Fractions containing purified Gal3–His were pooled, snap-frozen in liquid nitrogen, and stored at −80 °C. Protein concentration was determined by measuring sample absorbance at 280 nm using the molecular weight and the extinction coefficient calculated from the amino acid sequence (26 kDa and 35870 M^-1^cm^-1^).

Purified Gal3-His was fluorescently labeled using an NHS ester dye (e.g., Alexa Fluor 488) at a molar ratio of 1:4 (protein:dye). Briefly, the protein was diluted in PBS and buffered to pH 8.0 with sodium bicarbonate to support amine-reactive conjugation chemistry. 10% glycerol was included to stabilize the protein during labeling. The NHS dye was freshly prepared in anhydrous DMSO and immediately added to the protein solution. The reaction was incubated at RT for 2 h at 300 rpm on a thermomixer. After labeling, excess dye was removed by multiple rounds of buffer exchange using pre-equilibrated Zeba Spin Desalting Columns, 7 kDa cutoff. The labeled protein was then subjected to overnight dialysis at 4°C against PBS to ensure complete removal of unreacted dye and buffer equilibration. Following purification, the labeled Gal3 was aliquoted, snap-frozen in liquid nitrogen, and stored at −80 °C for long-term use.

#### GST-pull-down

GST-tagged proteins (0.1 µM) were immobilized onto GSH beads and incubated for 2 h at 4°C with 1.5 mg of MCF10A cellular lysate prepared in JS buffer (50 mM Hepes pH 7.5, 150 mM NaCl, 1% Triton X100, 5 mM EGTA, 1.5 mM MgCl_2_, 10% glycerol) supplemented with a protease inhibitor cocktail (Complete, EDTA-free, Roche). After three washes in JS buffer, bound proteins were eluted in 2X Laemmli buffer, resolved by SDS-PAGE and detected by IB. Samples were run on separate gels to detect proteins with similar molecular weights. Gels were then sectioned into the regions of interest to enable probing with different antibodies.

#### Whole-mount Carmine Alum staining of mammary glands

The inguinal (no. 4) mammary gland was dissected and fixed in 4% formalin overnight and preserved in 70% ethanol at 4°C. Fixed tissue was hydrated with ddH_2_O and stained in Carmine Alum overnight at RT. Post-staining, samples were dehydrated with a gradient of 70 – 95 – 100% ethanol. Fat tissues were clarified in a 1:2 solution of benzyl alcohol and benzyl benzoate (Sigma-Aldrich). Imaging was performed using a Stereomicroscope (Nikon SMZ-color camera Nikon DS-Fi3).

#### Mammary branching density analysis

Branching of the mammary glands was quantified from images of Carmine Alum-stained whole-mount mammary glands, as previously described in^15^. Briefly, starting from a determined area above the lymph node in all samples, we drew at least 100 linear intersections of 0.1 mm width per sample. We then quantified the number of branches crossing the 0.1 mm intersections using an automated macro in ImageJ software as in^15^. The mean number of all intersections for each mammary gland was reported as the mean branching density.

#### Mammary branching penetration analysis

Branching penetration of mammary glands was quantified from images of Carmine Alum-stained whole-mount mammary glands. The length of branched area was measured from the lymph node to the end of the branching, identified by the stained epithelial ducts. The full length of the mammary gland was measured from the lymph node to the end of the mammary fat pad, typically marked by the presence of muscle tissue at the edge of each whole mount. The mammary branching penetration percentage was calculated by the equation:

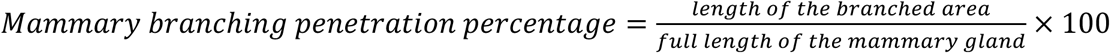

#### Immunohistochemistry on FFPE sections

Three-µm sections were prepared from FFPE blocks of inguinal mammary gland and dried at 37°C. The sections were processed with Bond-III fully Automated stainer system (Leica biosystems) according to the following protocol: tissue was deparaffinized and pre-treated with the Epitope Retrieval Solution 2 pH 9.0 (#AR9640) at 95°C for 20 min. After washing steps, peroxidase blocking was performed for 5 min using the Bond Polymer Refine Detection Kit (#DC9800, Leica Biosystems). Tissues were incubated for 30 min with rabbit polyclonal anti-EPN3 (in-house, 1:5000), rabbit monoclonal anti-c-erbB-2 (Dako #A0485, 1:1000) or rabbit monoclonal anti-Ki67 (Abcam #Ab16667, 1:50) diluted in Bond Primary Antibody Diluent (#AR9352). Subsequently, tissues were incubated with post primary and polymer solution for 8 min, developed with DAB-chromogen for 10 min, and counterstained with hematoxylin for 5 min. Ki67 positivity was calculated as the percentage of positive cells using QuPath software (v0.5.1).

#### Immunofluorescence on FFPE sections

Three-µm sections were prepared from FFPE blocks of inguinal mammary gland and dried at 37°C. The sections were deparaffinized, rehydrated through a graded ethanol series and pre-treated with 1 mM Tris-EDTA (pH 9.0) at 95°C for 50 min. After washing steps blocking was performed for 1 h at RT in blocking solution (TBS, 2% BSA, 2% donkey serum and 0.05% Tween). Tissues were incubated for 2 h at RT with rabbit polyclonal anti-EPN3 (in-house, 1:5000), rat monoclonal anti-KRT8 (clone TROMA-1 in-house, 1:200), or mouse monoclonal anti-KRT14 (Abcam #ab7800, 1:50). For IF, sections were incubated with secondary antibodies and nuclei were counterstained with DAPI. The images were acquired with NanoZoomer S80 Digital slide scanner (Hamamatsu, Japan) and assessed using QuPath software (v0.5.1).

#### Assessment of mammary cell subpopulations by FACS

Purified MECs were incubated with 0.2 U/ml Dispase® I (STEMCELL™ technologies) for 10 min at 37°C. Samples were then centrifuged at 335 g for 5 min at RT. Epithelial cells were purified using the EasySep™ Mouse Epithelial Cell Enrichment Kit (STEMCELL™ Technologies), according to the manufacturer’s protocol. Following purification, epithelial cells were incubated with 10% BSA-PBS for blocking and stained for 45 min at 4°C with primary fluorochrome-conjugated antibodies in 1% BSA-PBS against specific markers of interest (see Table S5). Stained cells were then fixed in 1% PFA and analyzed by FACS. Data acquisition and analysis were conducted using FlowJo software (v10.10.0). Cells were gated according to EpCAM-APC and Cd49f (eFluo450) levels. The proportion of positive cells for each marker was determined and normalized against single-stained controls to ensure accurate interpretation of marker expression levels.

#### ScRNA-seq and data analysis

After single cell digestion and mouse epithelial cell enrichment as described previously, cells were stained with live/dead viability stain (BV-510, BDenriched en) and EpCAM-APC antibody for 45 min at 4°C. Single, live EpCAM^pos^ cells were then purified by FACS-sorting using a BD InFlux V7 flow cytometer, and mixed with the reverse transcription mix using 10x Genomics Chromium Single-cell 3’ reagent kit (protocol V3.1) according to the manufacturer’s protocol and then loaded onto 10x Genomics Single-Cell 3’ Chips (www.10xgenomics.com). Libraries were sequenced using the Illumina Novaseq 6000 Sequencing System (with dual indexing format according to the manufacturer’s protocol V3.1) with a coverage of 50K reads per cell. Read alignment, UMI counting, cell-containing barcode (CB) calling were performed using the “cellranger count” from CellRanger (v6.0) on the mouse reference genome Mmu38. The different samples were aggregated using the “cellranger aggr” function, normalizing the depth across the input libraries. Subsequent analyses (quality filtering, normalization, dimensionality reduction, clustering and differential gene expression) were performed using Seurat v5.0.1^16^. Briefly, the pipeline included: i) filtering out low-quality cells with nCount_RNA ≥ 1000 and <60000, nFeature_RNA ≥ 200 and < median(nFeature_RNA) + 5 × MAD(nFeature_RNA) (MAD = median absolute deviation), and ≤ 10% mitochondrial genes; ii) Log normalization of the data with the “LogNormalize” normalization method; iii) variable feature identification with vst method and nfeatures=2000; iv) cell clustering on scaled data using as parameters dims = 10, k = 20 and r = 0.5; v) UMAP reduction^17^ with RunUMAP function; vi) specific markers of each cluster were identified using FindAllMarkers, pct.1 > 0.7 and top 10 were selected based on the avg_log2FC on merged clusters.

DEGs of each MEC subpopulation were identified using the FindMarkers function, with default parameters (pct = 0.01 and logfc = 0.1) comparing CTRL *vs.* EPN3-KI within main clusters (BM, LP and LH) and considering only non-proliferating cells (as assessed by proliferation score, Fig.S10A and Table S2). Functional analyses of BM, LP and LH clusters were generated with mSigDB using the upregulated genes in the EPN3-KI condition compared to CTRL (pvaladj < 0.05, avg_log2FC > 0.5). Signatures were then obtained using the UCell package (v2.6.2)^18^. Gene lists for lactation and alveologenesis signatures were obtained from previously published papers and manually curated as described in^19, 20^.

#### Transwell Matrigel invasion assay

Invasion assays were performed using 24-well Transwell™ Matrigel® Invasion Chambers with 8.0 µm PET membrane plates (Corning®). Primary MECs (30 x 10^3^/sample) were resuspended in serum-free medium and seeded in the upper chamber of the Transwell insert. The lower chamber contained 10% NA-FBS serum as a chemoattractant, while the negative control wells contained serum-free medium. The Transwell Invasion Chambers were incubated at 37°C for 24 h to allow cell invasion in the presence of I3 (20 µM), (+)-Lactose (100 mM) or vehicle DMSO. Then, the upper chamber was cleaned with a cotton swab and cells were fixed with 4% PFA for 10-15 min at RT. Cells were then washed in PBS + glycine 100 mM and stained with Hoechst (1 µg/ml). Invading cells were acquired with a Nikon Eclipse Ti2 microscope. At least 4 fields of view were captured from each technical replicate, and two technical replicates were considered in each experiment. Results were reported as the number of nuclei counted with ImageJ software (version 2.14.0).

#### Isolation of primary mammary organoids

To isolate primary organoids, mice were sacrificed, inguinal and thoracic mammary glands were excised and subjected to 1 h of enzymatic digestion in medium DMEM-Ham’s F12 medium (Gibco, Life Technologies), 3 mg/ml Collagenase IA (Sigma), 1.5 mg/ml Trypsin, 10 mM HEPES, and 1% Pen/Strep. Samples were incubated at 37°C in a humidified atmosphere containing 5% CO_2_ on rotation, 15 rpm.

The suspension was resuspended in PBS and centrifuged at 450 g for 10 min at RT. The upper layer containing stromal pieces was aspirated and the pellet was resuspended in PBS. If necessary, the suspension was incubated with 10 U/ml DNAse I (Merck Life science) for 2 min at RT followed by centrifugation at 450 g for 5 min. The supernatant was removed, and red blood cells were removed with ACK Lysing buffer for 1 min (Gibco). Samples were then centrifuged at 450 g for 5 min to obtain a pellet of mammary pieces. Mammary pieces were immediately resuspended in Matrigel (Corning) and plated in non-treated tissue culture plates (Corning) with MEGM medium (Lonza) for 5-7 days at 37°C in a humidified atmosphere containing 5% CO_2_, to form organoids. Organoids were detached with cold PBS and used for further analyses (see Branching morphogenesis assay and Organoid outgrowth).

#### Branching morphogenesis assay

Organoids were plated for branching morphogenesis following a previously defined protocol ^21^. Briefly, organoids were resuspended in 3:7 mix of neutralized Rat collagen I (∼3-4 mg/ml, Corning): Matrigel (Corning) and plated in ibiTreat-µ-Slide (Ibidi, 8-Well chambers) at density of ∼150-200 organoids/30 µl drop. After drop solidification, Basic Organoid Medium [BOM; DMEM-Ham’s F12 medium (Gibco, Life Technologies), ITS (Gibco)] supplemented with 2.5 nM FGF2 (PeproTech) and 3 μM Y-27632 (Rock inhibitor, PeproTech) was added for 24 h. Then, BOM supplemented with alternation of 2.5 nM FGF2 (PeproTech) or 2 nM EGF (PeproTech) was added every 2-3 days for a total of 10-14 days, with or without inhibitors: 100 mM (+)-Lactose, 20 μM I3, 4 µM Genz or DMSO. The images were obtained with Thunder microscope (Leica, DMI8) and the morphology of branched organoids was assessed at the endpoint of experiment.

#### Organoid outgrowth

Organoids were plated at ∼150-200 organoids/30 µl drop density in non-tissue culture treated plates (Corning) with MEGM medium (Lonza) supplemented with 3 μM Y-27632 (Rock inhibitor, PeproTech) for 24 h and then w/o rock-inhibitor for further outgrowth with inhibitors: 100 mM (+)-Lactose, 20 μM I3, 4 µM Genz or DMSO (for this specific assay, the 100 mM (+)-Lactose solution was prepared directly in MEGM isosmotic medium). Organoids were grown until sufficient material was obtained for RNA extraction (6-12 days).

#### RNA extraction and RT-qPCR analysis

To collect RNA, organoids were lysed in QIAzol™ Lysis Reagent (Qiagen) and total RNA was extracted with miRNeasy® micro kit (Qiagen) according to the manufacturer’s protocol. Total RNA was quantified by Qubit RNA HS assay (Invitrogen), RT-qPCR was performed using the SuperScript™ VILO™ cDNA synthesis kit (Invitrogen) and SsoAdvanced Universal Probes Supermix (Bio-Rad) on a LightCycler® 480 real-time PCR. The RT-qPCR Taqman primers used are listed below:

Gapdh: mm_9999915

Tbp: Mm01277042_g1

Actb: Mm00607939_s1

Csn1s1: Mm01160593_m1

Lalba: Mm00495258_m1

Data were normalized to the average expression levels of the housekeeping (HK) genes (*Gapdh*, *Tbp*, *Actb*) unless specified otherwise and results were presented as relative mRNA expression compared to HK genes, utilizing the 2^ΔCt method.

#### RNA extraction from FFPE lung blocks and RT-qPCR for metastases detection

Lungs from 28-30 week-old mice were included in FFPE blocks and 50 sections of 5 µm were pooled for RNA extraction. For each lung, the two halves were processed separately and considered as independent samples for RNA extraction. Briefly, the FFPE tissue sections were deparaffinized using xylene and physically disrupted by TissueLyser II (Qiagen) and Tungsten carbide beads for 5 min at 30 Hz. Samples were centrifuged for 4 min, full speed at RT followed by removal of supernatant. The xylene excess was removed using absolute ethanol and centrifugation for 4 min, full speed at RT followed by removal of supernatant. The ethanol excess was evaporated from the tissue pellet by spinning samples with an open lid using a SpeedVac™ for 8 min at 45°C. Then, the sample pellet was subjected to protein digestion by proteinase K in PKD buffer, agitating on a thermoblock for 15 min at 56°C and 450 rpm and immediate cooling down on ice for 3 min. The solution was centrifuged for 15 min, full speed at RT and the supernatant was processed for RNA isolation.

Total RNA was extracted with the AllPrep® DNA/RNA FFPE Kit (Qiagen) following manufacturer’s instructions, quantified by NanoDrop and reverse transcribed withSuperScript™ VILO™ cDNA Synthesis Kit (Invitrogen) according to the manufacturer’s instructions. cDNA was then pre-amplified using the TaqMan™ PreAmp Master Mix (Life Technologies) targeting genes of interest (10 amplification cycles) and used for quantitative PCR (qPCR) with gene-specific TaqMan™ assays with SsoAdvanced Universal Probes Supermix (Bio-Rad) on a LightCycler® 480 real-time PCR instrument. The RT-qPCR Taqman primers used are listed below:

Gapdh: mm_9999915

Tbp: mm01277042_g1

Gusb: mm01197698_m1

Erbb2: Rn01461176_g1

Erbb2: mm01306784_g1

Lalba: Mm00495258_m1

Data analysis involved calculating the cycle threshold (Ct) values. Relative gene expression was determined using the 2^-ΔCt method, with normalization to the average expression of three HK reference genes (*Gapdh, GusB*, and *Tbp)*. The qPCR assays were performed in three technical replicates to ensure accuracy and reproducibility.

## Statistics and software

All the analyses and statistics related were produced using Prism Software V10.2, JMP 18 (SAS) software or Microsoft Excel. Microsoft Excel or Prism were used to generate bar graphs. All the images were analyzed and quantified with ImageJ 1.54p.

**Fig. S1.**
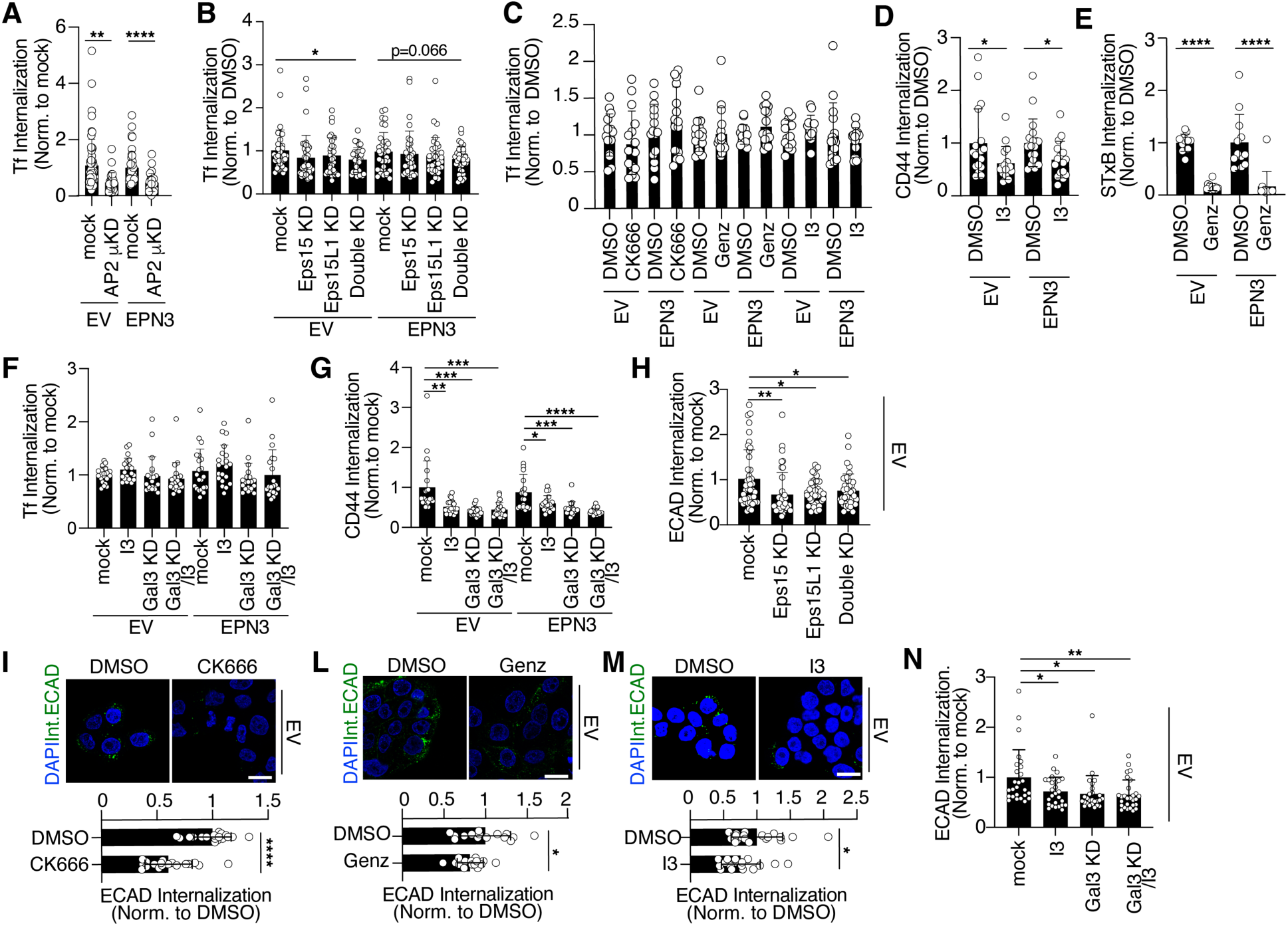
ECAD endocytosis in MCF10A cells is GL-Lect driven. **A)** Internalization of Tf-488 was monitored in MCF10A-EV and -EPN3 cells with or without AP2µ KD. Results are shown normalized to mock control. N (fields of view): EV, Mock=56, AP2µ KD=28 (n=3); EPN3, Mock=44, AP2µ KD=28 (n=3). **B)** Quantification of Tf-488 internalization in MCF10A-EV and - EPN3 cells subjected to single or double Eps15/Eps15L1 KDs. N (fields of view): EV, Mock=35, Eps15 KD=35, Eps15L1 KD=35, Double KD=35 (n=3); EPN3, Mock=36, Eps15 KD=35, Eps15L1 KD=35, Double KD=35 (n=3). **C)** Negative controls for inhibitor screening (supporting Fig. 1D). Quantification of Tf-488 internalization in MCF10A-EV and -EPN3 cells pre-treated with the following compounds or vehicle control: CK666 (50 µM, 1 h), Genz (4 µM, 6 days), I3 (20 µM, 10 min). N (fields of view): EV, DMSO=15, CK666=15, Genz=15, I3=15 (n=3); EPN3, DMSO=15, CK666=15, Genz=15, I3=14 (n=3). **D-E)** Positive controls for inhibitor screening (supporting Fig. 1D). (D) CD44 internalization was monitored *in vivo* by IF using an anti-CD44 antibody in MCF10A-EV and -EPN3 cells pre-treated with I3 (20 µM, 10 min) or vehicle control. Quantification of relative CD44 fluorescence intensity is shown normalized to control. N (fields of view): EV, DMSO=18, I3=16; EPN3, DMSO=18, I3=16 (n=3). (E) Shiga-toxin (STXB) endocytosis and binding was monitored by continuous incubation of STXB-488 conjugated ligand in MCF10A-EV and- EPN3 cells pre-treated with Genz (4 µM, 6 days). Quantification of relative STXB fluorescence intensity is shown normalized to control. N (fields of view): EV, DMSO=12, Genz=12; EPN3, DMSO=12, Genz=12 (n=3). **F)** Quantification of Transferrin internalization in MCF10A-EV and -EPN3 cells subjected to I3 (20 µM, 10 min) treatment, Gal3 KD or Gal3 KD/I3 treatment. N (fields of view): EV, mock=20, I3=20, Gal3 KD=20, Gal3 KD/I3=20 (n=2); EPN3, mock=21, I3=20, Gal3 KD=21, Gal3 KD/I3=21 (n=2). **G)** Quantification of CD44 internalization in MCF10A-EV and -EPN3 cells subjected to I3 (20 µM, 10 min) treatment, Gal3 KD or Gal3 KD/I3 treatment. N (fields of view): EV, mock=20, I3=20, Gal3 KD=21, Gal3 KD/I3=20 (n=2); EPN3, mock=20, I3=19, Gal3 KD=20, Gal3 KD/I3=20 (n=2). **H)** ECAD internalization was monitored in MCF10A-EV cells subjected to Eps15/Eps15L1 single or double KD. Quantification of relative ECAD fluorescence intensity/cell normalized to mock control. N (fields of view): Mock=42, Eps15 KD=39, Eps15L1 KD=37 and Double KD=40 (n=3). **I-M)** ECAD internalization was monitored in MCF10A-EV cells pre-treated as in panel C. Top, representative images showing internalized ECAD (green) and DAPI staining (blue). Bar, 20 µm. Bottom, quantification of relative ECAD fluorescence intensity is shown normalized to control. N (fields of view): (I) DMSO=18, CK666=18; (L) DMSO=15, Genz=15; (M) DMSO=19, I3=17 (n=3). **N)** ECAD internalization was monitored in MCF10A-EV cells subjected to I3 (20 µM, 10 min) treatment, Gal3 KD or Gal3 KD/I3 treatment. N (fields of view): EV, mock=26, I3=27, Gal3 KD=26, Gal3 KD/I3=28 (n=2). p-values in the relevant panels (Unpaired Student’s t-test, two-tailed): ****, <0.0001; ** <0.001, * <0.05

**Fig. S2.**
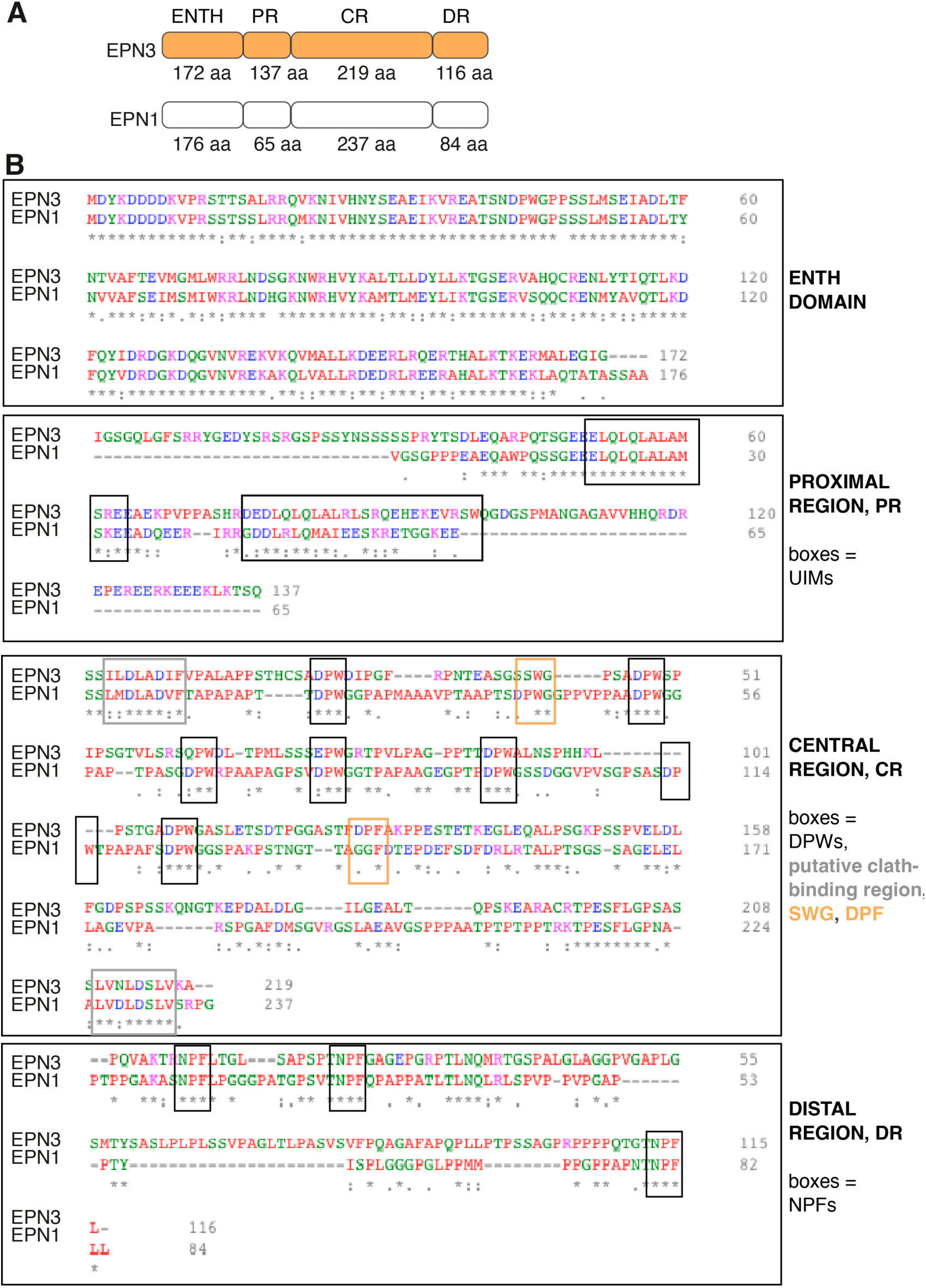
EPN3 and EPN1 protein domain organization. **A)** Schematic representation of the four domains of EPN3 and EPN1 proteins which were swapped to generate chimeras: epsin-N-terminal homology domain (ENTH), proximal region (PR), central region (CR) and distal region (DR). Size of the domains in indicated by number of amino acids (aa). **B)** Clustal Omega alignments of the amino acid sequences of each EPN3 and EPN1 domain. Ubiquitin interacting motifs (UIMs) are shown and boxed in the PR; DPWs, putative clathrin binding regions, SWG and DPF motifs are boxed in the CR; NPF motifs are boxed in the DR.

**Fig S3.**
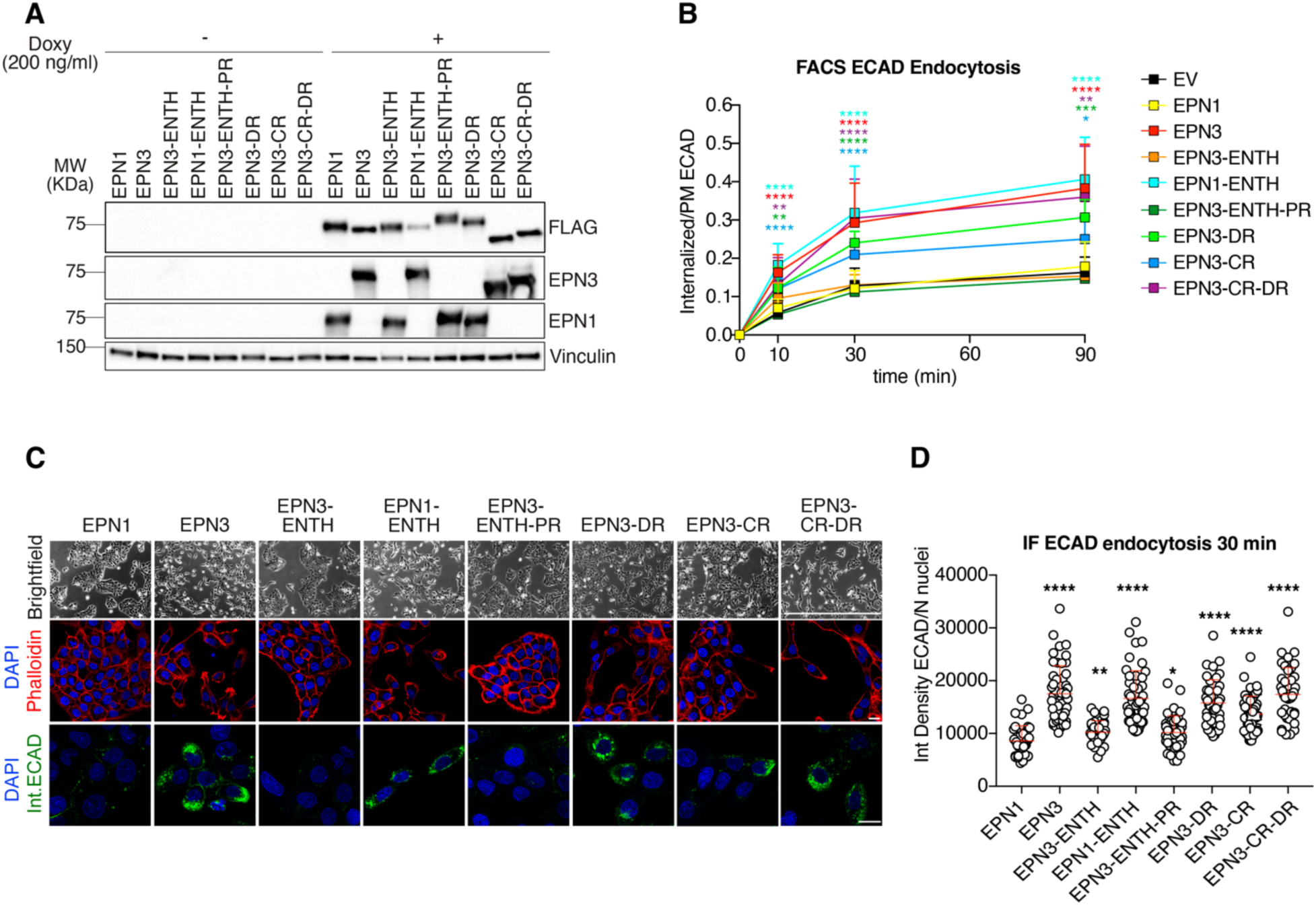
Molecular determinants mediating EPN3-dependent phenotypes are localized in the central and distal regions. **A)** Lysates of MCF10A cells expressing the indicated doxy-inducible FLAG-tagged EPN3, EPN1, chimeras or EV control (induced or not with 200 ng/ml doxy for 24h) were subjected to immunoblotting with home-made EPN1 or EPN3 antibodies. Vinculin, loading control. Molecular weights (MWs) are indicated on left. In KDa. **B)** Time-course of ECAD internalization measured by FACS. Data are reported as mean fluorescence intensity of internalized ECAD signal over the total plasma membrane (PM) signal ± SD (n≥3). **C)** Top, representative brightfield microscopy images of MCF10A cells described in A. Bar, 1000 μm. Middle, representative confocal images of phalloidin staining which was used to measure the aspect ratio of each cell. Red Phalloidin, blue DAPI. Bar, 20 µm. Bottom, representative confocal images of ECAD internalization at 30 min measured by IF. Green, internalized ECAD, blue DAPI. Bar, 20 µm. **D)** Quantification of ECAD internalization at 30 min measured by IF, supporting the results obtained by FACS. Results are reported as integrated density of fluorescence intensity divided by the number of nuclei in that field of view. N of nuclei>200. p-values in relevant panels (Unpaired Student’s t-test two-tailed) *vs* EPN1; ****, <0.0001; ***, <0.001, **, <0.01; * <0.05.

**Fig S4.**
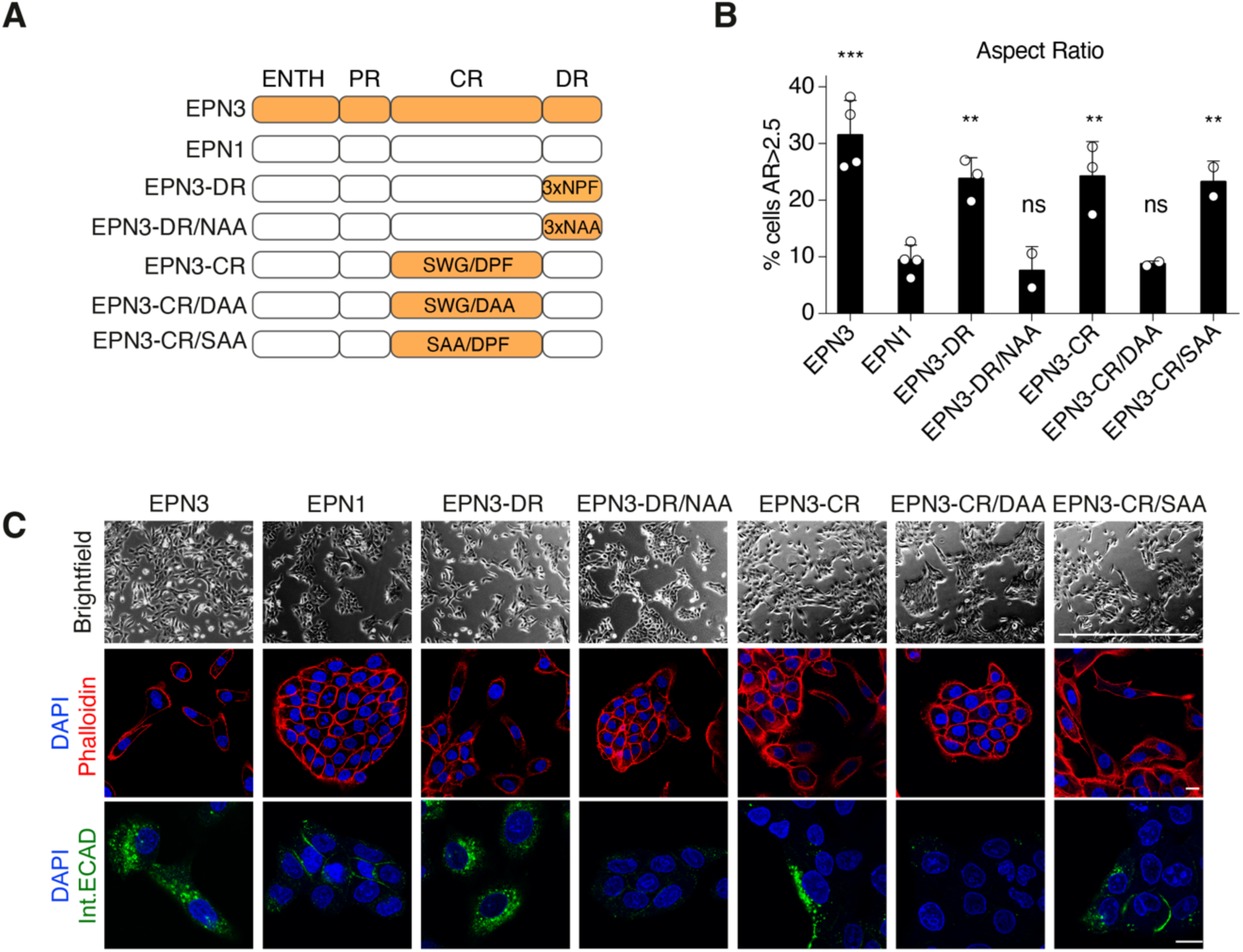
Mutagenesis of Eps15/Eps15L1 binding sites abrogate EPN3-dependent phenotypes. **A)** Schematic representation of the FLAG-tagged EPN3 and EPN1 WT and mutant chimeras: EPN3-DR with 3 WT NPFs compared to EPN3-DR/NAA in which all 3 NPF motifs were mutagenized into NAA; EPN3-CR with WT SWG/DPF motifs compared to EPN3-CR/DAA in which the DPF motif is mutagenized into DAA and EPN3-CR/SAA where the SWG motif is mutagenized into SAA. **B)** Phalloidin staining of cells expressing the constructs shown in A was used to measure the aspect ratio of each cell. Percentage of cells with an aspect ratio > 2.5 is shown ± SD; n≥2. p-value (Unpaired Student’s t-test two-tailed) *vs.* EPN1; ***, <0.001, **, <0.01; ns, not significant. **C)** Top, representative brightfield images of MCF10A cells expressing the constructs described in A. Bar, 1000 μm. Middle, representative confocal images of phalloidin staining used to measure the aspect ratio of each cell shown in B. Red Phalloidin, blue DAPI. Bar, 20 µm. Bottom, representative confocal images of ECAD internalization at 90 min measured by IF. Green ECAD, blue DAPI. Bar, 20 µm.

**Figure S5.**
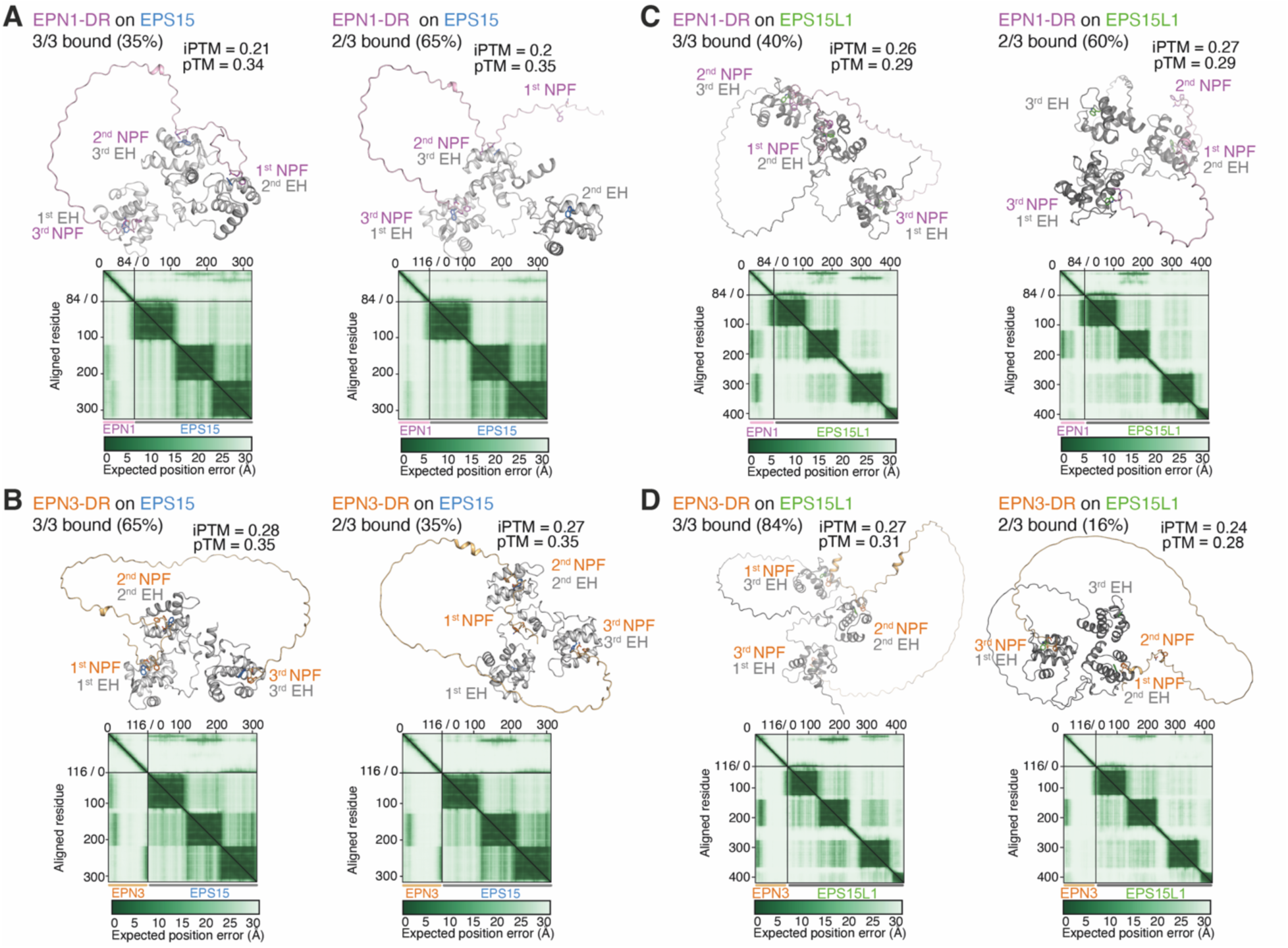
AlphaFold structural modeling of EPN3-DR and EPN1-DR binding to the EH domain-containing region of Eps15 and Eps15L1. AlphaFold3 modelling was used to predict the complexes formed between the EPN1/EPN3-DR regions and the Eps15/Eps15L1 EH domains. All possible models, relative frequencies represented as percentages, interface predicted template modelling (iPTM) and predicted template modelling (pTM) scores, and predicted aligned error (PAE) plots are shown (n=50). In particular, models are shown for: **A)** EPN1-DR and Eps15 EH domains; **B)** EPN3-DR and Eps15 EH domains; **C)** EPN1-DR and Eps15L1 EH domains; **D)** EPN3-DR and Eps15L1 EH domains. Of note, AF3 predictions displayed an overall low ipTM (<0.3), likely due to the long unstructured loops that characterize the DR domains. Nevertheless, at least two out of three NPF residues in the EPN1/EPN3-DR domains were predicted to bind to the EH domain with good PAE scores, in the same binding pocket that harbors the aromatic residue previously identified as critical for ligand interaction^22^. This prompted us to perform multiple rounds of AF3 modeling with different starting seeds to compare the frequency of interactions between EPN1/EPN3 NPF motifs and the EH domains of Eps15/Eps15L1. In all generated models, the third NPF site exhibited a higher binding frequency in EPN3 compared to EPN1 (65% for EPN3 *vs.* 35% for EPN1 with Eps15; 84% for EPN3 *vs.* 40% for EPN1 with EPS15L1).

**Figure S6.**
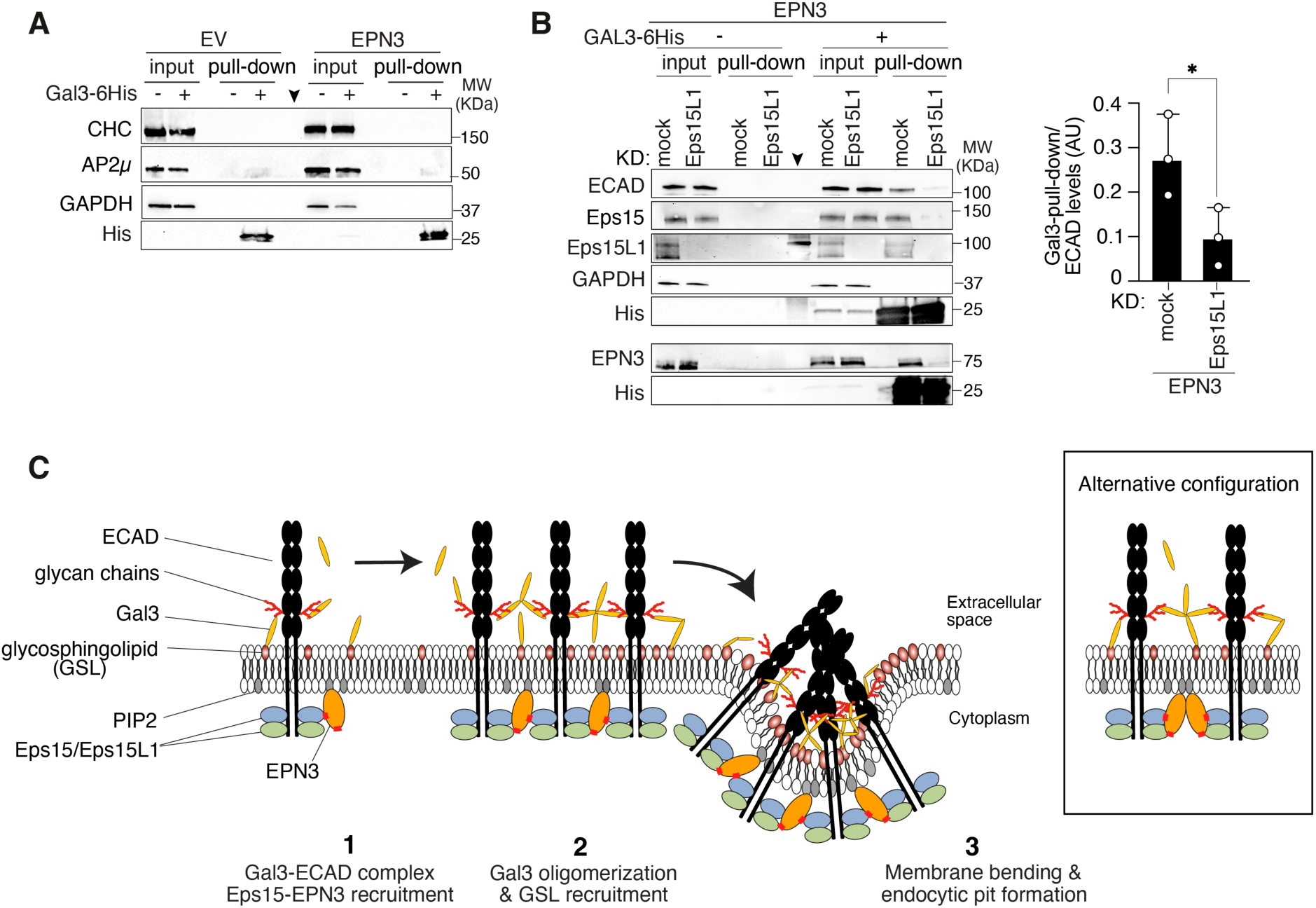
EPN3 coordinates an Eps15/Eps15L1-mediated GL-Lect mechanism driving ECAD endocytosis. **A)** Additional controls for Gal3 pull-down shown in Fig. 3A. MCF10A-EV or -EPN3 cells were incubated or not with recombinant human Gal3-6His for 1 h on ice, lysed and subjected to pull-down with His-Fab trap. IB was as shown on the left. GAPDH is used as negative control for pull-down and loading control for inputs. MWs are indicated on the right in KDa. A vertical arrow marks the lane loaded with the molecular MW marker. **B)** Eps1L1 KD affects ECAD-EPN3 recruitment to Gal3. MCF10A-EPN3 cells were transfected with siRNAs targeting Eps15L1 (or mock). Then, cells were incubated or not with recombinant human Gal3-6His for 1 h on ice, lysed and subjected to pull-down with His-Fab trap. IB was as shown on the left; GAPDH is used as negative control for pull-down and loading control for inputs. MWs are indicated on the right in KDa. A vertical arrow marks the lane loaded with the molecular MW marker. Right, quantification of ECAD protein levels, normalized to His levels in the pull-down; n=3. p-value (Unpaired Student’s t-test two-tailed) *vs* EV: * <0.05. **C)** Cartoon representing a working model of EPN3-mediated GL-Lect-driven ECAD endocytosis. The model is based on the following information: i) EPN3 is required for ECAD endocytosis [REF^1^ and Fig. 2A]. ii) EPN3 does not directly interact with ECAD; in contrast, Eps15 and Eps15L1 both associate with ECAD (Fig. 3C). The structural interfaces responsible for these interactions remain undefined, and it is not yet established whether the interactions are direct or mediated by intermediary proteins. iii) Eps15 and Eps15L1 perform non-redundant functions in ECAD internalization (Fig. 1B). Given that Eps15/Eps15L1 heterodimers have been experimentally demonstrated^23, 24^, these findings suggest that such heterodimers represent the functional unit mediating ECAD endocytosis. iv) In further support of this notion, single KD of either Eps15 or Eps15L1 markedly impaired the recruitment of ECAD and EPN3 to His-Gal3 (Fig. 3B, and **Fig. S6B**), and the recruitment of one adaptor in the Gal3 pull-down was affected by depletion of the other. v) EPN3 interacts with both Eps15 and Eps15L1 with comparable efficiency *in vivo* (Fig. 2D). Two distinct binding interfaces on EPN3 mediate these interactions: one located within the central CR region, where a DPF motif plays a pivotal role, and another in the C-terminal DR region, which contains NPF motifs (Fig. 2E, H). Both DPF and NPF motifs are recognized binding determinants for the EH domains found in the N-termini of Eps15 and Eps15L1. Notably, these interaction surfaces are either absent in EPN1 (*e.g.,* the DPF motif; **Fig. S2**) or display reduced binding efficiency toward Eps15/Eps15L1 (*e.g.,* the NPF motifs, as shown in FLAG-coimmunoprecipitation *in vivo* (Fig. 2D) and as predicted by AlphaFold modeling; Fig. 2G **and S5**). In the schematic representation, ECAD (black) forms dimers at the plasma membrane (PM)^25,26^. On the cytoplasmic side, Eps15/Eps15L1 heterodimers (blue and green) associate with the ECAD intracellular domain, thereby linking ECAD to EPN3 (orange). EPN3, in turn, is anchored to the inner leaflet of the PM via its ENTH domain, which interacts with phosphatidylinositol 4,5-bisphosphate (PIP₂, gray ovals)^27^. Through its dual interaction surfaces (highlighted in red), EPN3 facilitates ECAD clustering by bridging two Eps15/Eps15L1 heterodimers associated with distinct ECAD molecules. This clustering promotes the recruitment of Galectin-3 (Gal3, yellow ovals) to N-glycosylated ECAD residues (red branches)^28, 29^. Oligomerized Gal3 (potentially forming tetramers; Shafaq-Zadah M. *et al.*, Nat Comm*, in press*) further bridges ECAD and glycosphingolipids (GSLs, red ovals), thereby organizing PM nanodomains that promote membrane curvature and endocytic pit formation. Additionally, EPN3’s binding to PIP₂ (gray ovals) on the cytosolic leaflet may enhance membrane remodeling and curvature generation. Alternative configurations are also conceivable (right inset). For instance, EPN3 may exist as a dimer^30^, potentially bridging two Eps15/Eps15L1 complexes through its dual interaction domains, thereby further stabilizing the endocytic complex.

**Figure S7.**
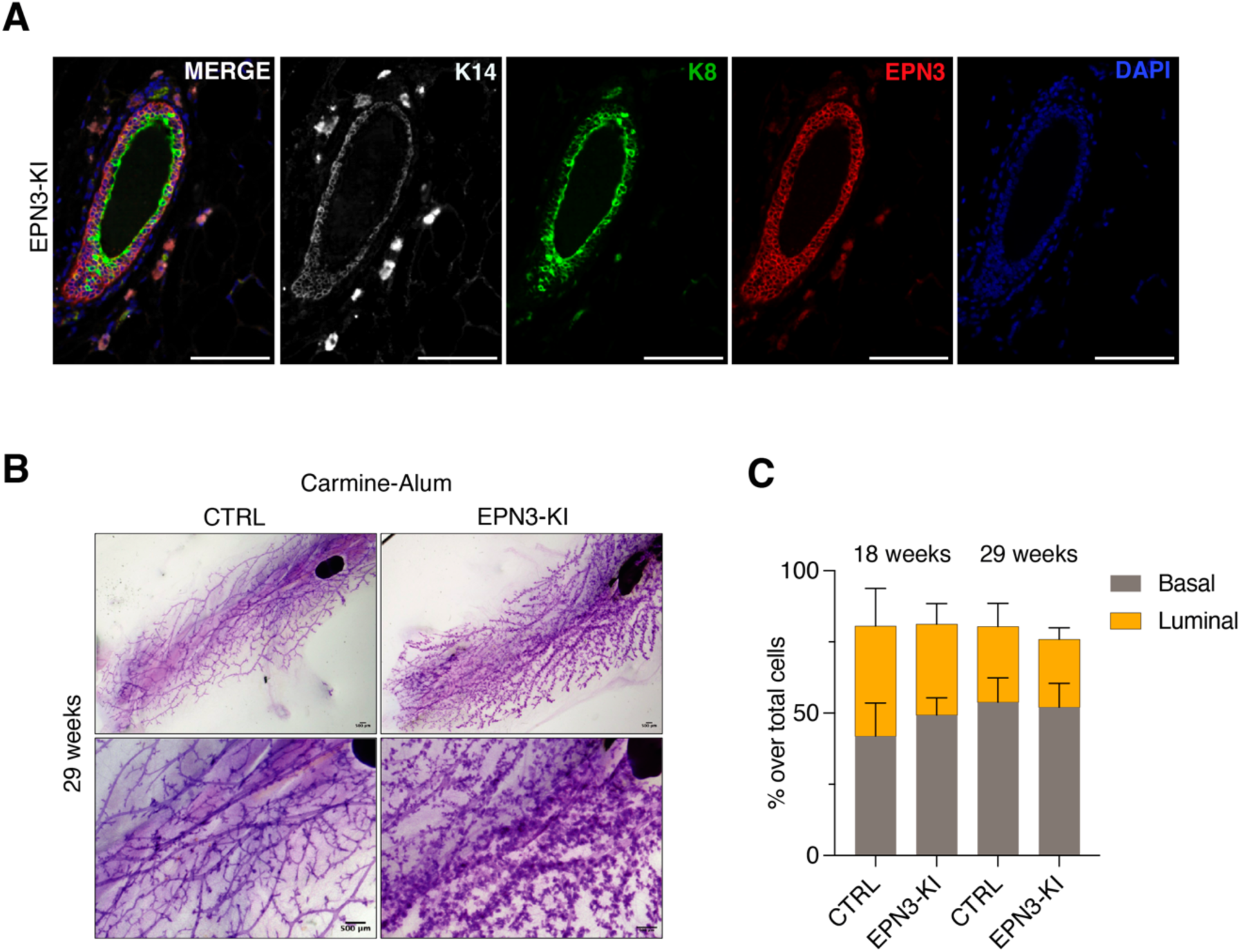
Characterization of EPN3-KI mice at 18 and 29 weeks of age. **A)** IF analysis of EPN3 expression in the mammary glands of EPN3-KI mice. Merged image shows localization of EPN3 expression (red) in an epithelial duct and its co-localization with luminal (K8, green) and basal (K14, white) markers, followed by single color images of K14, K8, EPN3 and DAPI within the same field of view, Bars, 100 μm. **B)** Representative images of whole-mount sections of mammary glands from 29-week-old CTRL (N=15) and EPN3-KI (N=14) mice; magnifications in lower panels: Bars, 500 μm. **C)** FACS analysis of basal and luminal mammary epithelial cell subpopulations at 18 weeks (CTRL, N=5; EPN3-KI, N=7) and 29 weeks (N=8 per condition). The percentage of each subpopulation over the total cells (basal and luminal, with exclusion of stromal cells) was calculated based on EpCAM and Cd49f expression evaluated by FACS analysis.

**Figure S8.**
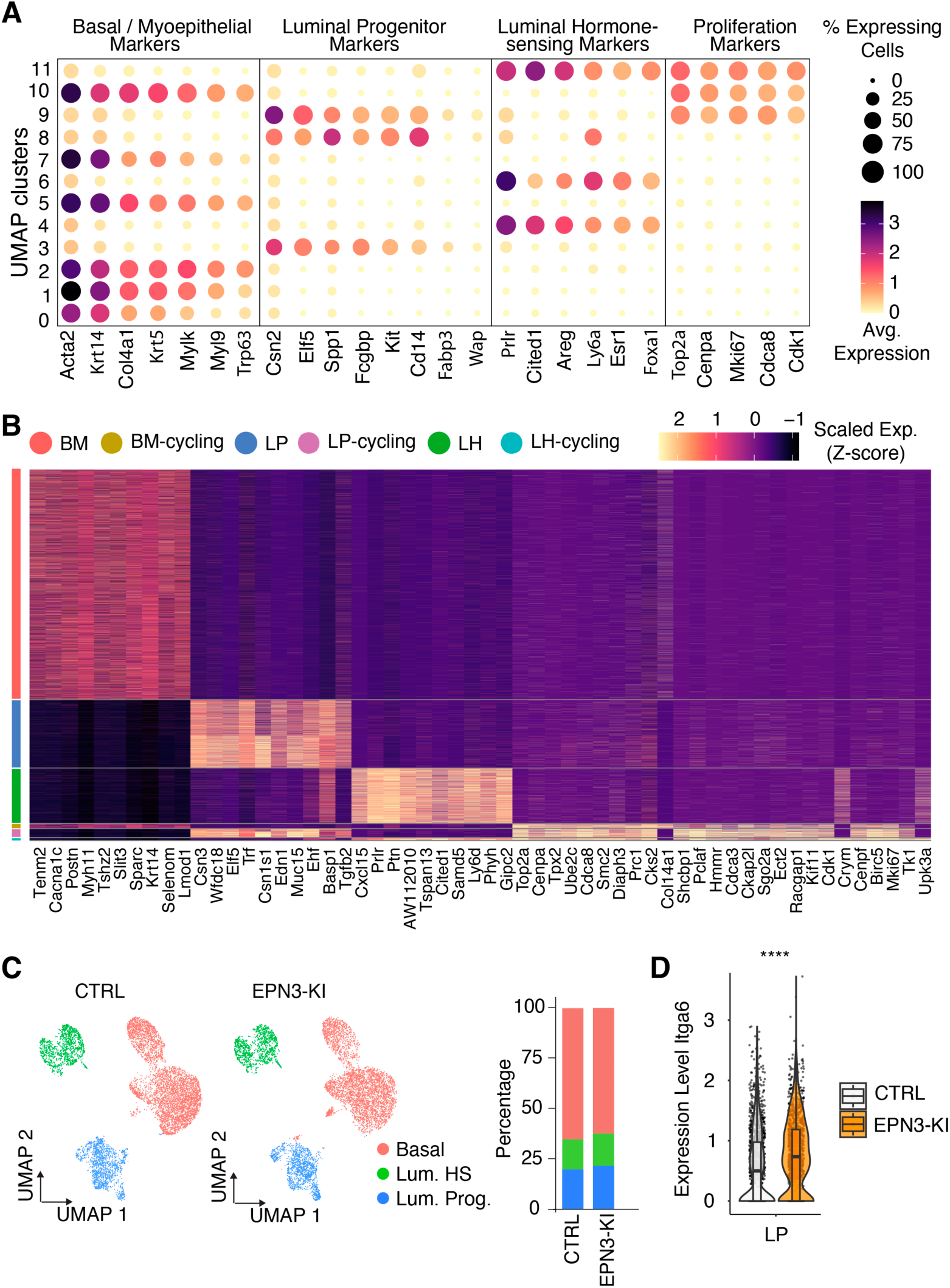
scRNA-seq analysis of EPN3-KI *vs*. CTRL mammary epithelial subpopulations. **A)** Dot plot expression of basal/myoepithelial (BM), luminal progenitor (LP), luminal hormone-sensing (LH) and proliferation markers. The color gradient shows the expression level from low (yellow) to high (black) and the dot size represents the percentage of expressing cells in merged UMAPs for each cluster depicted in **Fig 5D**. **B)** Heatmap representation showing the differentially expressed genes per main cell cluster (BM, LP, and LH) and their proliferating clusters (BM-Cycling, LP-Cycling, and LH-Cycling). Each cluster is color-coded, and the size of each cluster reflects its relative abundance, calculated as the proportion of cells within that cluster compared to the total number of cells analyzed. The color gradient from black to yellow represents gene expression levels, with black indicating low expression and yellow indicating high expression. The x-axis lists the top 10 upregulated genes in each cluster. **C)** Left, unsupervised UMAP plot showing color-coded mammary epithelial subpopulations (BM, LP, LH) in both CTRL and EPN3-KI samples. Right, stacked bar graph showing the percentage of each subpopulation over the total number of cells. **D)** Violin plot shows the expression levels of Itga6 in the CTRL and EPN3-KI LP populations. Statistical analysis performed with the Wilcoxon rank sum test and Bonferroni correction, comparing each cluster in EPN3-KI *vs.* CTRL, **** <0.0001.

**Figure S9.**
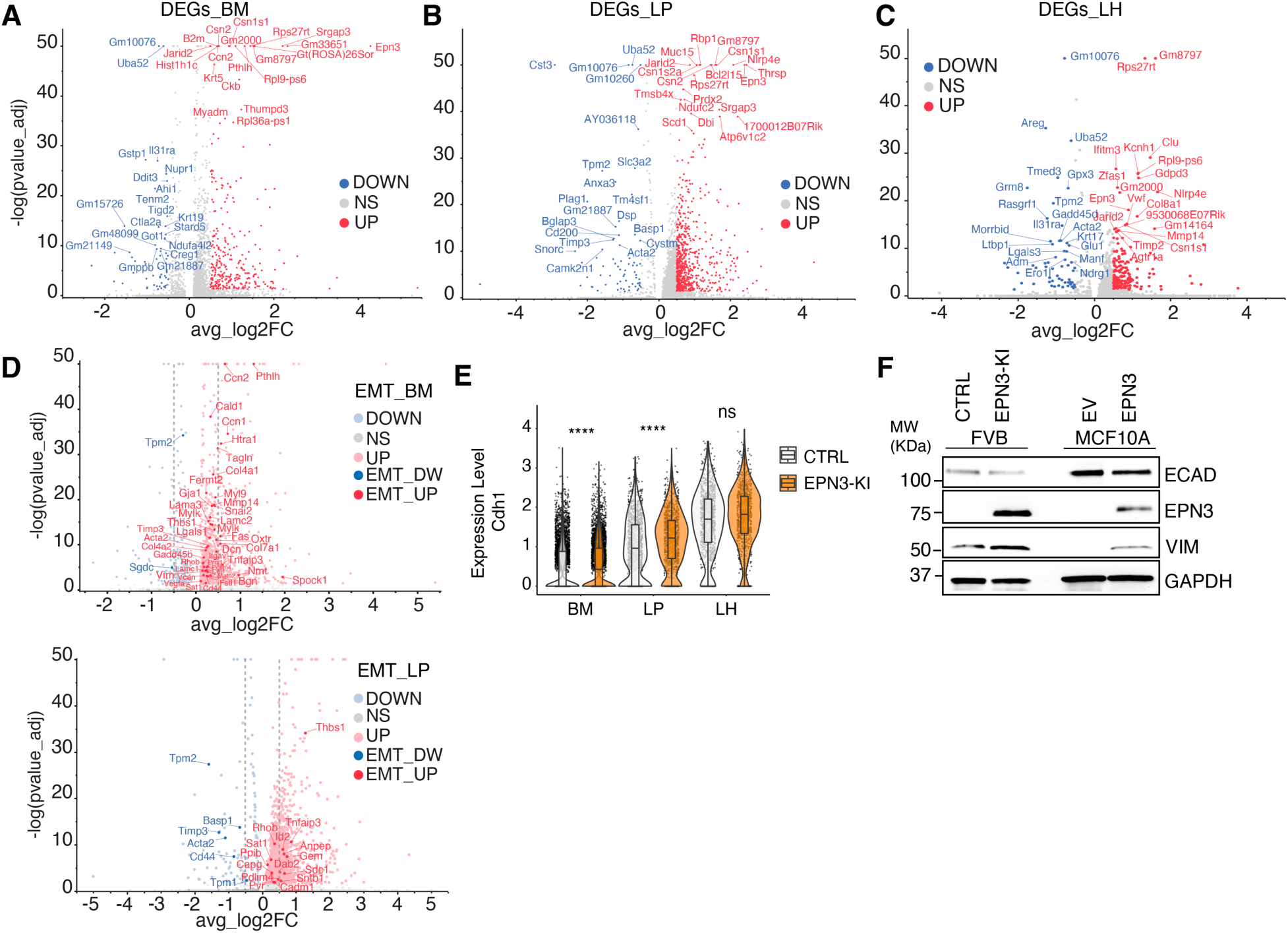
scRNA seq analysis of differentially expressed genes in EPN3-KI *vs*. CTRL mammary epithelial subpopulations. **A-C)** Volcano plots of differentially expressed genes (DEGs) in basal/myoepithelial (BM) (**A**), luminal progenitor (LP) (**B**), luminal hormone sensing (LH) (**C**) mammary epithelial subpopulations derived from EPN3-KI and CTRL mice. Log2 fold-change (avg_log2FC) of >|0.5| and adjusted p-value (p-value_adj) <0.05. Downregulated (DOWN); upregulated (UP); no significant difference (NS). Top 20 significantly upregulated or downregulated genes are highlighted in each panel. For graphical reasons, genes with -log(p-value_adj) > 50, were capped at 50. **D)** Volcano plots of EMT genes (from EMT_Hallmark collection, mSigDb) in basal/myoepithelial (EMT_BM, top) and luminal progenitor (EMT_LP, bottom) regulated in EPN3-KI *vs*. CTRL. In light red/blue are shown all the DEGs with log2 fold-change (avg_log2FC) of >|0.1| and adjusted p-value (p-value_adj) < 0.05. Downregulated (DOWN); upregulated (UP); no significant difference (NS). EMT genes significantly upregulated (dark red, EMT_UP) or downregulated (EMT_DW) are highlighted in each panel. For graphical reasons, those genes with -log(p-value_adj) > 50, were capped at 50. Dotted lines highlight the threshold of (avg_log2FC) of >|0.5| and adjusted p-value (p-value_adj) < 0.05, used in panels A-C. **E**) Violin plot showing the expression levels of the ECAD gene *Cdh1* in the three populations. Statistical analysis was performed with Wilcoxon rank sum test and Bonferroni correction, comparing each cluster in the EPN3-KI *vs.* CTRL samples, **** p_val_adj <0.0001, ns, not significant. **F**) IB analysis of ECAD expression in mammary epithelial cells from CTRL and EPN3-KI mice, or in MCF10A-EV and -EPN3 cells. GAPDH was used as a protein loading control. MWs are shown on the left in KDa.

**Figure S10.**
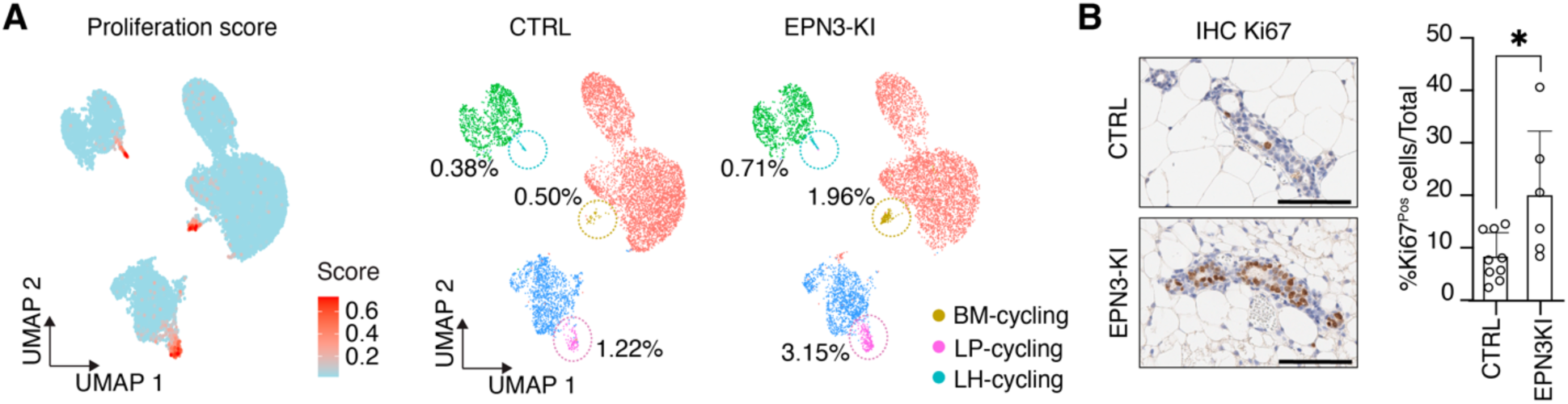
scRNA-seq analysis of proliferating cells within EPN3-KI and CTRL mammary epithelial subpopulations. **A)** Left, UMAP plot showing the proliferation score, measured in scRNA-seq profiles based on the expression levels of 5 markers (**Table S2**). The quantification of actively proliferating cells in each cluster (% over the total number of cells), marked by circles, is shown in separate UMAP plots for CTRL (middle panel) and EPN3-KI (right panel) MECs. **B)** Left, representative IHC images of Ki67 expression in the mammary glands of CTRL and EPN3-KI mice at 18 weeks (bar, 100 μm). Right, the percentage of Ki67-positive cells over total number of cells in each slide analyzed was determined; mean % + SD is reported (CTRL=9, EPN3-KI= 6). p-value (Unpaired Student’s t-test, two-tailed); * <0.05.

**Figure S11.**
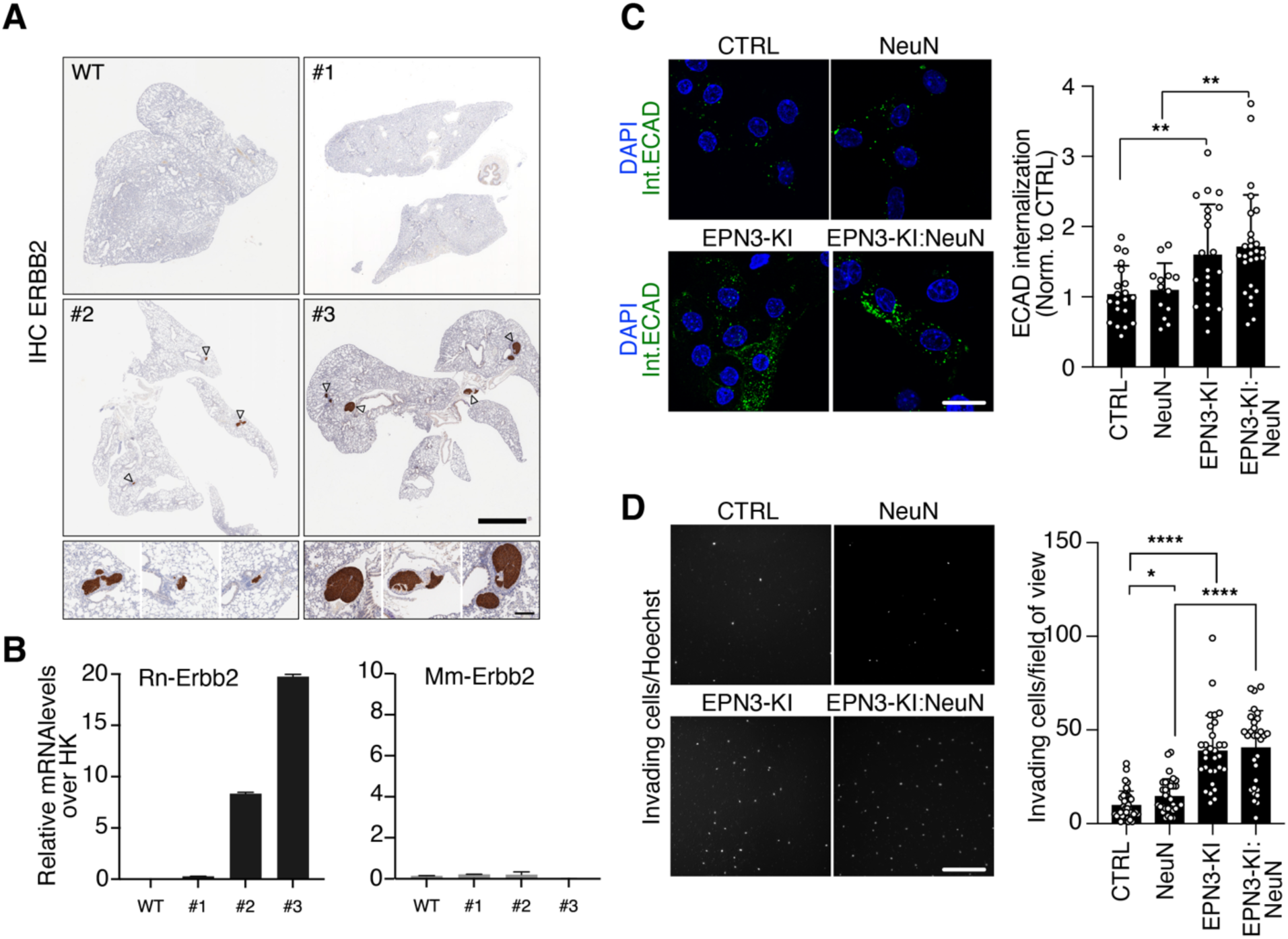
EPN3-dependent ECAD endocytosis and invasiveness is active in a tumorigenic context. **A)** Representative anti-ERBB2 IHC staining of lung tissue samples from CTRL and EPN3-KI:MMTV-NeuN mice (#1 to #3) with varying metastatic loads (from the same sample cohort used in Fig. 7E). Bar, 2 mm. Arrowheads point to metastatic lesions magnified in insets, bar, 200 µm. **B)** RT-qPCR analysis of endogenous Mm-Erbb2 and transgenic Rn-Erbb2 (NeuN) in lung tissues collected in (A) and shown as relative levels over HK genes. Standard deviation is calculated from technical qPCR triplicate. Samples were used to validate the RT-qPCR assay for metastasis detection shown in Fig. 7E. **C)** ECAD internalization was monitored by IF at 37°C for 30 min in primary MECs derived from CTRL (N=3), NeuN (N=4), EPN3-KI (N=4) and EPN3-KI:MMTV-NeuN mice (N=4). Left, representative IF images are shown: internalized ECAD (green), DAPI (blue). Bar, 20 µm. Right, quantification of ECAD fluorescence intensity normalized to control, n= 3. p-values (Unpaired Student’s t-test, two-tailed): **, <0.01; ns, not significant. **D)** Transwell Matrigel invasion assay. Left, representative fields of view of invading cell nuclei stained with Hoechst; bar, 200 µm. Right, quantification of invading cells expressed as the number of invading cells per field of view; (the plot contains the samples used for generating panels Fig. 6C and Fig. 7H). N (mice): (CTRL, N=9; EPN3-KI, N=8; NeuN, N=8; EPN3-KI:MMTV-NeuN, N=7). N (fields of view): CTRL=40, NeuN=30, EPN3-KI=40, and EPN3-KI:MMTV-NeuN=30. p-values (Unpaired Student’s t-test, two-tailed): ****, <0.0001; ns, not significant.

**Table S1. Additional data related to Figures 1-3. A) Pharmacological and genetic perturbations of endocytic mechanisms and their effects on ECAD and control cargo uptake.** The table, from left to right, lists the treatments, target molecule(s), endocytic mechanism(s) primarily affected, and the fraction of internalization over control cells (mean ± SEM) for ECAD, CD44, STxB, Dextran, and Transferrin (Tf). Target molecules involved in ECAD internalization are highlighted in red, ECAD is the primary cargo investigated in this study. CD44 uptake serves as a marker for GL-Lect–dependent endocytosis, Shiga toxin B subunit (STxB) reflects glycosphingolipid-dependent uptake, Dextran reports macropinocytosis, and Tf is the canonical marker for clathrin-mediated endocytosis (CME). Inhibition of GL-Lect pathways (Lactose, I3, Genz) reduces both ECAD and CD44 uptake, whereas macropinocytosis inhibition (EIPA) selectively impairs Dextran internalization. Inhibition of actin polymerization (CK666) decreases ECAD internalization but not Tf, while AP2 knockdown (AP2μ-KD) selectively reduces Tf uptake, confirming CME specificity. Caveolin KD and RTN3 KD showed no significant effects under the tested conditions, both on ECAD and Tf. “nd” indicates not determined, and “ns” denotes changes that did not reach statistical significance. Reported p values were calculated using an unpaired Student’s t-test *vs*. untreated control. **B) Validation of target knockdowns in MCF10A cells.** Representative immunoblots (IB) validating the KD of the indicated target proteins in MCF10A-EV and -EPN3 cells. KDs were performed using siRNA at 8 nM over two consecutive cycles (5 days in total) to ensure robust suppression of protein expression. Shown are representative blots for: i) EPS15, EPS15L1, and the double KD of EPS15/EPS15L1, which participate in both CME and NCE; ii) AP2µ in both MCF10A-EV and - EPN3, which specifically regulates CME; iii) RTN3 in both MCF10A-EV and -EPN3, which primarily targets NCE; and iv) Caveolin 1/2 double KD in MCF10A-EPN3 cells only, assessing caveolae-mediated endocytosis. v) Gal3 KD in MCF10A-EV and -EPN3 cells, assessing Gal3-mediated GL-Lect driven endocytosis. These blots confirm the efficiency and specificity of each KD summarized in the **Table S1A**.

**Table S2. Additional data related to Fig. 5. Gene_Signatures.** List of genes used for calculating the different signature scores in the scRNA-seq profiles. **Mammary epithelial cell (MEC) subpopulations identified by scRNA-seq.** Total cell counts and relative percentages of MEC subpopulations in CTRL and EPN3-KI samples obtained from scRNA-seq data. Percentages were calculated relative to the total number of cells in each respective sample The ratio indicates the relative abundance of each subpopulation in EPN3-KI compared to CTRL samples.

**Table S3. Differentially expressed genes (DEGs) in EPN3 KI.** Differential expression analysis comparing EPN3-KI to CTRL in basal/myoepithelial (BM), luminal progenitor (LP), and luminal hormone-sensing (LH) subpopulations (excluding actively proliferating clusters). Each sheet, reports genes significantly upregulated (DEGs_UP, avg_log2FC>0.5, p_val_adj<0.05) or downregulated (DEGs_DOWN, avg_log2FC< -0.5, p_val_adj< 0.05) within each subpopulation. Columns include: gene_name (gene symbol), avg_log2FC (average log2 fold-change of EPN3-KI vs CTRL), pct.1 (percentage of cells expressing the gene in EPN3-KI), pct.2 (percentage of cells expressing the gene in CTRL), p_val (Wilcoxon rank-sum test, default in the Seurat FindMarkers function), and p_val_adj (adjusted P-value).

**Table S4. Pathways Enriched in scRNAseq profiles.** Pathway enrichment analysis of upregulated DEGs in BM, LP, and LH subpopulations from scRNA-seq data. For each subpopulation, Hallmark gene sets are listed together with the gene set name and its description, number of genes included in the gene set, number of genes in overlap, k/k, p-value, FDR-q-value and -log10[FDR q-value] are listed as “Overlap Results”.

In addition, tables listing genes belonging to each Hallmark gene set (white columns), alongside the overlapping genes between the upregulated DEGs and the corresponding Hallmark set are shown for each subpopulation as “Pathways”.

**Table S5. List of Reagents**. The tables summarize the list of antibodies (Abs), tissue-culture (TC) reagents, and primers used to generate chimeric EPN1- and EPN3-constructs employed in this study.

## References

1. Nieto, M.A., Huang, R.Y., Jackson, R.A. & Thiery, J.P. Emt: 2016. Cell 166, 21–45 (2016).

2. Youssef, K.K. & Nieto, M.A. Epithelial-mesenchymal transition in tissue repair and degeneration. Nat Rev Mol Cell Biol 25, 720–739 (2024).

3. Pastushenko, I. & Blanpain, C. EMT Transition States during Tumor Progression and Metastasis. Trends Cell Biol (2018).

4. Hu, M. et al. Regulation of in situ to invasive breast carcinoma transition. Cancer Cell 13, 394–406 (2008).

5. Yang, J. et al. Guidelines and definitions for research on epithelial-mesenchymal transition. Nat Rev Mol Cell Biol 21, 341–352 (2020).

6. Jolly, M.K., Mani, S.A. & Levine, H. Hybrid epithelial/mesenchymal phenotype(s): The ‘fittest’ for metastasis? Biochim Biophys Acta Rev Cancer 1870, 151–157 (2018).

7. Grigore, A.D., Jolly, M.K., Jia, D., Farach-Carson, M.C. & Levine, H. Tumor Budding: The Name is EMT. Partial EMT. J Clin Med 5 (2016).

8. Aiello, N.M. et al. EMT Subtype Influences Epithelial Plasticity and Mode of Cell Migration. Dev Cell 45, 681–695 e684 (2018).

9. Lanzetti, L. & Di Fiore, P.P. Behind the Scenes: Endo/Exocytosis in the Acquisition of Metastatic Traits. Cancer Res 77, 1813–1817 (2017).

10. Schiano Lomoriello, I., et al. A self-sustaining endocytic-based loop promotes breast cancer plasticity leading to aggressiveness and pro-metastatic behavior. Nat Commun 11, 3020 (2020).

11. Horvath, C.A., Vanden Broeck, D., Boulet, G.A., Bogers, J. & De Wolf, M.J. Epsin: inducing membrane curvature. Int J Biochem Cell Biol 39, 1765–1770 (2007).

12. Ko, G. et al. Selective high-level expression of epsin 3 in gastric parietal cells, where it is localized at endocytic sites of apical canaliculi. Proc Natl Acad Sci U S A 107, 21511–21516 (2010).

13. Wood, K.M. & Smith, C.J. Clathrin: the molecular shape shifter. Biochem J 478, 3099–3123 (2021).

14. Pascolutti, R. et al. Molecularly Distinct Clathrin-Coated Pits Differentially Impact EGFR Fate and Signaling. Cell Rep 27, 3049–3061 e3046 (2019).

15. Sigismund, S. et al. Clathrin-independent endocytosis of ubiquitinated cargos. Proc Natl Acad Sci U S A 102, 2760–2765 (2005).

16. Howes, M.T. et al. Clathrin-independent carriers form a high capacity endocytic sorting system at the leading edge of migrating cells. J Cell Biol 190, 675–691 (2010).

17. Peterson, K. et al. Systematic Tuning of Fluoro-galectin-3 Interactions Provides Thiodigalactoside Derivatives with Single-Digit nM Affinity and High Selectivity. J Med Chem 61, 1164–1175 (2018).

18. Kay, R.R. Macropinocytosis: Biology and mechanisms. Cells Dev 168, 203713 (2021).

19. Parton, R.G., Tillu, V.A. & Collins, B.M. Caveolae. Curr Biol 28, R402–R405 (2018).

20. Caldieri, G. et al. Reticulon 3-dependent ER-PM contact sites control EGFR nonclathrin endocytosis. Science 356, 617–624 (2017).

21. de Beco, S., Amblard, F. & Coscoy, S. New insights into the regulation of E-cadherin distribution by endocytosis. Int Rev Cell Mol Biol 295, 63–108 (2012).

22. Johannes, L., Shafaq-Zadah, M., Dransart, E., Wunder, C. & Leffler, H. Endocytic Roles of Glycans on Proteins and Lipids. Cold Spring Harb Perspect Biol 16 (2024).

23. Polo, S., Confalonieri, S., Salcini, A.E. & Di Fiore, P.P. EH and UIM: endocytosis and more. Sci STKE 2003, re17 (2003).

24. Krieger, J.R. et al. Identification and selected reaction monitoring (SRM) quantification of endocytosis factors associated with Numb. Mol Cell Proteomics 12, 499–514 (2013).

25. Paoluzi, S. et al. Recognition specificity of individual EH domains of mammals and yeast. EMBO J 17, 6541–6550 (1998).

26. Santonico, E., Panni, S., Falconi, M., Castagnoli, L. & Cesareni, G. Binding to DPF-motif by the POB1 EH domain is responsible for POB1-Eps15 interaction. BMC Biochem 8, 29 (2007).

27. Coda, L. et al. Eps15R is a tyrosine kinase substrate with characteristics of a docking protein possibly involved in coated pits-mediated internalization. J Biol Chem 273, 3003–3012 (1998).

28. Cupers, P., ter Haar, E., Boll, W. & Kirchhausen, T. Parallel dimers and anti-parallel tetramers formed by epidermal growth factor receptor pathway substrate clone 15. J Biol Chem 272, 33430–33434 (1997).

29. Tebar, F., Confalonieri, S., Carter, R.E., Di Fiore, P.P. & Sorkin, A. Eps15 is constitutively oligomerized due to homophilic interaction of its coiled-coil region. J Biol Chem 272, 15413–15418 (1997).

30. Fu, N.Y., Nolan, E., Lindeman, G.J. & Visvader, J.E. Stem Cells and the Differentiation Hierarchy in Mammary Gland Development. Physiol Rev 100, 489–523 (2020).

31. Scheele, C.L. et al. Identity and dynamics of mammary stem cells during branching morphogenesis. Nature 542, 313–317 (2017).

32. Virtanen, S., Schulte, R., Stingl, J., Caldas, C. & Shehata, M. High-throughput surface marker screen on primary human breast tissues reveals further cellular heterogeneity. Breast Cancer Res 23, 66 (2021).

33. Paavolainen, O. & Peuhu, E. Integrin-mediated adhesion and mechanosensing in the mammary gland. Semin Cell Dev Biol 114, 113–125 (2021).

34. Xu, R., Boudreau, A. & Bissell, M.J. Tissue architecture and function: dynamic reciprocity via extra- and intra-cellular matrices. Cancer Metastasis Rev 28, 167–176 (2009).

35. Bach, K. et al. Differentiation dynamics of mammary epithelial cells revealed by single-cell RNA sequencing. Nat Commun 8, 2128 (2017).

36. Langille, E. et al. Loss of Epigenetic Regulation Disrupts Lineage Integrity, Induces Aberrant Alveogenesis, and Promotes Breast Cancer. Cancer Discov 12, 2930–2953 (2022).

37. Pervolarakis, N. et al. Integrated Single-Cell Transcriptomics and Chromatin Accessibility Analysis Reveals Regulators of Mammary Epithelial Cell Identity. Cell Rep 33, 108273 (2020).

38. Caruso, M., Huang, S., Mourao, L. & Scheele, C. A Mammary Organoid Model to Study Branching Morphogenesis. Front Physiol 13, 826107 (2022).

39. Guy, C.T. et al. Expression of the neu protooncogene in the mammary epithelium of transgenic mice induces metastatic disease. Proc Natl Acad Sci U S A 89, 10578–10582 (1992).

40. Muller, W.J. et al. Synergistic interaction of the Neu proto-oncogene product and transforming growth factor alpha in the mammary epithelium of transgenic mice. Mol Cell Biol 16, 5726–5736 (1996).

41. Mukherjee, S., Louie, S.G., Campbell, M., Esserman, L. & Shyamala, G. Ductal growth is impeded in mammary glands of C-neu transgenic mice. Oncogene 19, 5982–5987 (2000).

42. Shyamala, G., Chou, Y.C., Cardiff, R.D. & Vargis, E. Effect of c-neu/ErbB2 expression levels on estrogen receptor alpha-dependent proliferation in mammary epithelial cells: implications for breast cancer biology. Cancer Res 66, 10391–10398 (2006).

43. Harper, K.L. et al. Mechanism of early dissemination and metastasis in Her2(+) mammary cancer. Nature 540, 588–592 (2016).

44. Hosseini, H. et al. Early dissemination seeds metastasis in breast cancer. Nature 540, 552–558 (2016).

45. Zhao, Z.M. et al. Early and multiple origins of metastatic lineages within primary tumors. Proc Natl Acad Sci U S A 113, 2140–2145 (2016).

46. Johannes, L., Jacob, R. & Leffler, H. Galectins at a glance. J Cell Sci 131 (2018).

47. Banfer, S. et al. Molecular mechanism to recruit galectin-3 into multivesicular bodies for polarized exosomal secretion. Proc Natl Acad Sci U S A 115, E4396–E4405 (2018).

48. Lakshminarayan, R. et al. Galectin-3 drives glycosphingolipid-dependent biogenesis of clathrin-independent carriers. Nat Cell Biol 16, 595–606 (2014).

49. Shafaq-Zadah, M. et al. Exploration into Galectin-3 Driven Endocytosis and Lattices. Biomolecules 14 (2024).

50. Lepur, A., Salomonsson, E., Nilsson, U.J. & Leffler, H. Ligand induced galectin-3 protein self-association. J Biol Chem 287, 21751–21756 (2012).

51. Howes, M.T., Mayor, S. & Parton, R.G. Molecules, mechanisms, and cellular roles of clathrin-independent endocytosis. Curr Opin Cell Biol 22, 519–527 (2010).

52. Carvalho, S. et al. Preventing E-cadherin aberrant N-glycosylation at Asn-554 improves its critical function in gastric cancer. Oncogene 35, 1619–1631 (2016).

53. Pinho, S.S. et al. Modulation of E-cadherin function and dysfunction by N-glycosylation. Cell Mol Life Sci 68, 1011–1020 (2011).

54. Ewald, A.J., Brenot, A., Duong, M., Chan, B.S. & Werb, Z. Collective epithelial migration and cell rearrangements drive mammary branching morphogenesis. Dev Cell 14, 570–581 (2008).

55. Shamir, E.R. & Ewald, A.J. Adhesion in mammary development: novel roles for E-cadherin in individual and collective cell migration. Curr Top Dev Biol 112, 353–382 (2015).

56. Fata, J.E., Werb, Z. & Bissell, M.J. Regulation of mammary gland branching morphogenesis by the extracellular matrix and its remodeling enzymes. Breast Cancer Res 6, 1–11 (2004).

57. Inman, J.L., Robertson, C., Mott, J.D. & Bissell, M.J. Mammary gland development: cell fate specification, stem cells and the microenvironment. Development 142, 1028–1042 (2015).

58. Oakes, S.R., Hilton, H.N. & Ormandy, C.J. The alveolar switch: coordinating the proliferative cues and cell fate decisions that drive the formation of lobuloalveoli from ductal epithelium. Breast Cancer Res 8, 207 (2006).

59. Twigger, A.J. et al. Transcriptional changes in the mammary gland during lactation revealed by single cell sequencing of cells from human milk. Nat Commun 13, 562 (2022).

60. Valdes-Mora, F. et al. Single-cell transcriptomics reveals involution mimicry during the specification of the basal breast cancer subtype. Cell Rep 35, 108945 (2021).

## References

1. Schiano Lomoriello, I., et al. A self-sustaining endocytic-based loop promotes breast cancer plasticity leading to aggressiveness and pro-metastatic behavior. Nat Commun 11, 3020 (2020).

2. Schorpp, M. et al. The human ubiquitin C promoter directs high ubiquitous expression of transgenes in mice. Nucleic Acids Res 24, 1787–1788 (1996).

3. Ramirez, A. et al. A keratin K5Cre transgenic line appropriate for tissue-specific or generalized Cre-mediated recombination. Genesis 39, 52–57 (2004).

4. Guy, C.T. et al. Expression of the neu protooncogene in the mammary epithelium of transgenic mice induces metastatic disease. Proc Natl Acad Sci U S A 89, 10578–10582 (1992).

5. Muller, W.J. et al. Synergistic interaction of the Neu proto-oncogene product and transforming growth factor alpha in the mammary epithelium of transgenic mice. Mol Cell Biol 16, 5726–5736 (1996).

6. Sorme, P., Kahl-Knutsson, B., Huflejt, M., Nilsson, U.J. & Leffler, H. Fluorescence polarization as an analytical tool to evaluate galectin-ligand interactions. Anal Biochem 334, 36–47 (2004).

7. Sorme, P., Kahl-Knutson, B., Wellmar, U., Nilsson, U.J. & Leffler, H. Fluorescence polarization to study galectin-ligand interactions. Methods Enzymol 362, 504–512 (2003).

8. Lakshminarayan, R. et al. Galectin-3 drives glycosphingolipid-dependent biogenesis of clathrin-independent carriers. Nat Cell Biol 16, 595–606 (2014).

9. Aiello, N.M. et al. EMT Subtype Influences Epithelial Plasticity and Mode of Cell Migration. Dev Cell 45, 681–695 e684 (2018).

10. Elfmann, C. & Stulke, J. PAE viewer: a webserver for the interactive visualization of the predicted aligned error for multimer structure predictions and crosslinks. Nucleic Acids Res 51, W404–W410 (2023).

11. Caldieri, G. et al. Reticulon 3-dependent ER-PM contact sites control EGFR nonclathrin endocytosis. Science 356, 617–624 (2017).

12. Mesa, D. et al. A tripartite organelle platform links growth factor receptor signaling to mitochondrial metabolism. Nat Commun 15, 5119 (2024).

13. Shafaq-Zadah, M. et al. Exploration into Galectin-3 Driven Endocytosis and Lattices. Biomolecules 14 (2024).

14. Bolte, S. & Cordelieres, F.P. A guided tour into subcellular colocalization analysis in light microscopy. J Microsc 224, 213–232 (2006).

15. Stanko, J.P. & Fenton, S.E. Quantifying Branching Density in Rat Mammary Gland Whole-mounts Using the Sholl Analysis Method. J Vis Exp (2017).

16. Hao, Y. et al. Dictionary learning for integrative, multimodal and scalable single-cell analysis. Nat Biotechnol 42, 293–304 (2024).

17. Becht, E. et al. Dimensionality reduction for visualizing single-cell data using UMAP. Nat Biotechnol (2018).

18. Andreatta, M. & Carmona, S.J. UCell: Robust and scalable single-cell gene signature scoring. Comput Struct Biotechnol J 19, 3796–3798 (2021).

19. Valdes-Mora, F. et al. Single-cell transcriptomics reveals involution mimicry during the specification of the basal breast cancer subtype. Cell Rep 35, 108945 (2021).

20. Langille, E. et al. Loss of Epigenetic Regulation Disrupts Lineage Integrity, Induces Aberrant Alveogenesis, and Promotes Breast Cancer. Cancer Discov 12, 2930–2953 (2022).

21. Caruso, M., Huang, S., Mourao, L. & Scheele, C. A Mammary Organoid Model to Study Branching Morphogenesis. Front Physiol 13, 826107 (2022).

22. Confalonieri, S. & Di Fiore, P.P. The Eps15 homology (EH) domain. FEBS Lett 513, 24–29 (2002).

23. Cupers, P., ter Haar, E., Boll, W. & Kirchhausen, T. Parallel dimers and anti-parallel tetramers formed by epidermal growth factor receptor pathway substrate clone 15. J Biol Chem 272, 33430–33434 (1997).

24. Tebar, F., Confalonieri, S., Carter, R.E., Di Fiore, P.P. & Sorkin, A. Eps15 is constitutively oligomerized due to homophilic interaction of its coiled-coil region. J Biol Chem 272, 15413–15418 (1997).

25. Boggon, T.J. et al. C-cadherin ectodomain structure and implications for cell adhesion mechanisms. Science 296, 1308–1313 (2002).

26. Harrison, O.J. et al. The extracellular architecture of adherens junctions revealed by crystal structures of type I cadherins. Structure 19, 244–256 (2011).

27. Itoh, T. et al. Role of the ENTH domain in phosphatidylinositol-4,5-bisphosphate binding and endocytosis. Science 291, 1047–1051 (2001).

28. Carvalho, S. et al. Preventing E-cadherin aberrant N-glycosylation at Asn-554 improves its critical function in gastric cancer. Oncogene 35, 1619–1631 (2016).

29. Pinho, S.S. et al. Modulation of E-cadherin function and dysfunction by N-glycosylation. Cell Mol Life Sci 68, 1011–1020 (2011).

30. Lai, C.L. et al. Membrane binding and self-association of the epsin N-terminal homology domain. J Mol Biol 423, 800–817 (2012).

